# Behavioral features of motivated response to alcohol in *Drosophila*

**DOI:** 10.1101/2020.02.17.953026

**Authors:** Jamie L. Catalano, Nicholas Mei, Reza Azanchi, Sophia Song, Tyler Blackwater, Ulrike Heberlein, Karla R. Kaun

**Affiliations:** Molecular Pharmacology and Physiology Graduate Program, Brown University, Providence, RI, USA; Department of Neuroscience, Brown University, Providence, RI, USA; Howard Hughes Medical Institute, Janelia Research Campus (HHMI JRC), Ashburn, VA, USA

**Keywords:** *Drosophila*, behavior, addiction, alcohol, self-administration, learning, memory, operant, instrumental, runway

## Abstract

Animals avoid predators and find the best food and mates by learning from the consequences of their behavior. However, reinforcers are not always uniquely appetitive or aversive but can have complex properties. Most intoxicating substances fall within this category; provoking aversive sensory and physiological reactions while simultaneously inducing overwhelming appetitive properties. Here we describe the subtle behavioral features associated with continued seeking for alcohol despite aversive consequences. We developed an automated runway apparatus to measure how *Drosophila* respond to consecutive exposures of a volatilized substance. Behavior within this Behavioral Expression of Ethanol Reinforcement Runway (BEER Run) demonstrated a defined shift from aversive to appetitive responses to volatilized ethanol. Behavioral metrics attained by combining computer vision and machine learning methods, reveal that a subset of 9 classified behaviors and component behavioral features associate with this shift. We propose this combination of 9 behaviors can be used to navigate the complexities of operant learning to reveal motivated goal-seeking behavior.

## Introduction

Animals define value and adapt to their environment by learning the consequences of their behavior. In a natural environment, stimuli are complex, forcing animals to integrate appetitive and aversive properties to initiate appropriate behavioral responses. As an animal learns new information with consecutive experiences, it initiates an adaptive response. However, we have very little understanding of the micro-behaviors that define these subtle behavioral shifts. In depth understanding of these subtle behaviors is key to understanding how the brain processes complex stimuli.

Intoxicating drugs are powerful tools to study these subtle behavioral features. They have both aversive and appetitive properties, and ultimately induce lasting reward drives, which persist despite aversive consequences. Understanding the switch in response from aversion to reward will clarify how drugs misregulate the brain’s valence system to result in compulsive seeking behavior. To understand how the reward response overrides the aversive response, it’s critical to first understand the behavioral features associated with this switch.

The recent development of high-throughput tracking methods coupled with machine learning has drastically improved the resolution and capability of behavioral characterization. This consequently provides an opportunity to increase our understanding of motivated behavior to better inform neurobiological efforts. A runway model of self-administration is ideal for investigating this switch because it permits tight control of timing and amount of drug delivery, requires a goal-directed response to release the drug, and has adequate space to observe a range of behavioral features that change with consecutive trials. It also permits dissociation of the behavioral features associated with Classical conditioning, where an animal associates cues with the intoxicating experience, and Operant conditioning, where a behavioral response is reinforced by the drug.

Runway paradigms have been effectively used to study motivated behavior for cocaine [1-3], opioids [4-6], nicotine [7, 8] and alcohol [9] in rodent models. We developed a runway-based memory apparatus for *Drosophila*, the Behavioral Expression of Ethanol Reinforcement Runway (BEER Run), in which the consequence of traversing the runway and entering the mechanically gated end chamber is an intoxicating dose of volatilized ethanol. This assay coupled with high-throughput tracking and machine learning allows for behavioral characterization of drug-seeking motivated behavior.

We demonstrate *Drosophila* can effectively be used to identify subtle behavioral features that occur as a consequence of aversive and appetitive experiences. Through analysis of 25 behaviors, we found that the intersection between latency to leave the start chamber, average velocity and pausing defines the switch in preference indicative of motivation for alcohol. This thorough behavioral characterization coupled with the extensive array of neurogenetic tools available in *Drosophila* provides a high resolution, high precision opportunity to understand the neurobiological basis of motivated operant behavior.

## Results

The BEER Run is an automated runway platform (Figures 1A and Additional Supplementary Item 1A) that provides an opportunity to examine animal behavior during the preconditioning, conditioning and postconditioning phases of operant learning. We sought to 1) identify an ethanol concentration that sufficiently stimulated seeking behavior, 2) understand how behavior changes as animals learn a task, and 3) identify a subset of behaviors that can be accurately used to measure motivated operant behavior in animals.

**Figure 1.**
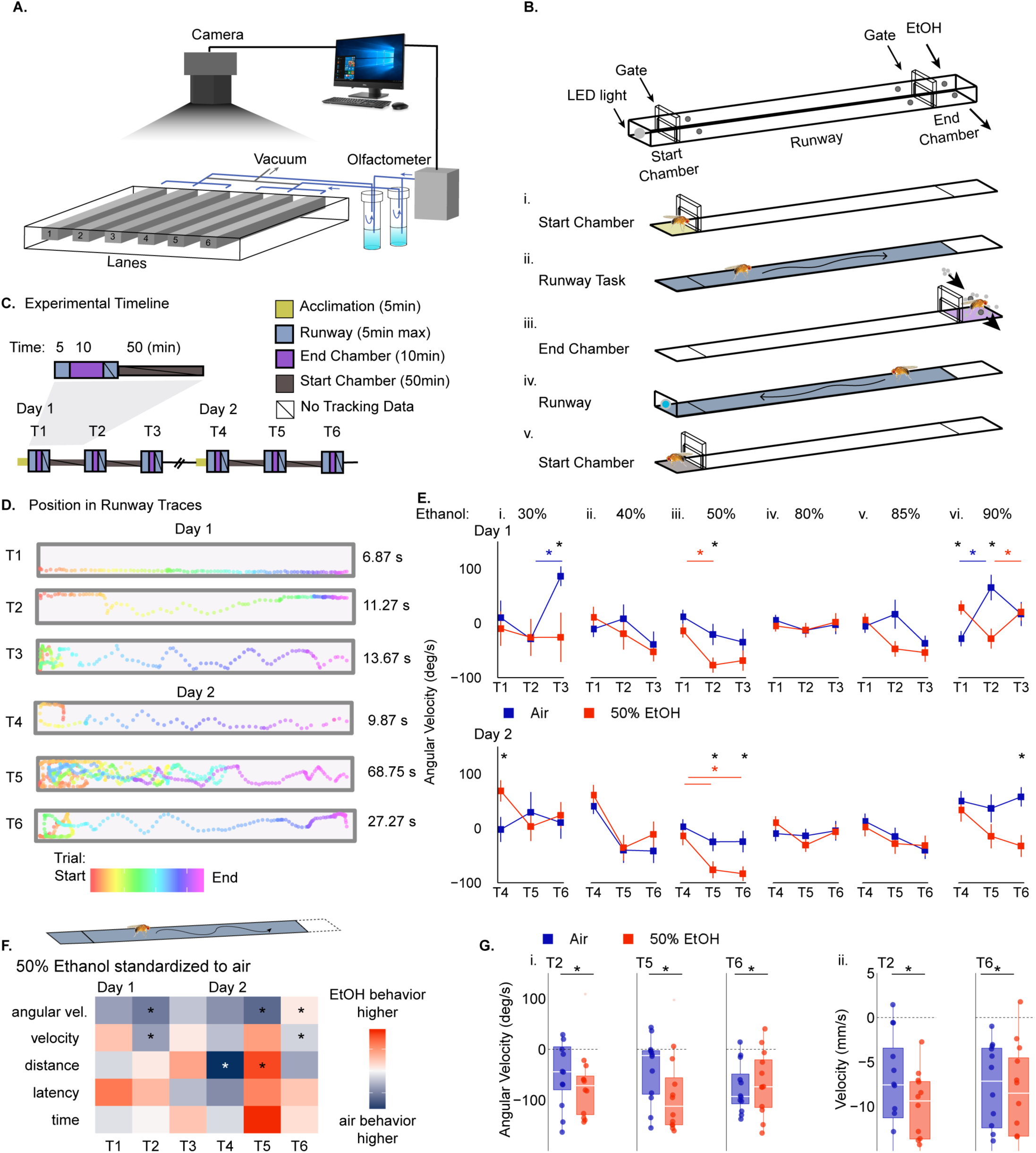
Behavioral expression of ethanol reinforcement Runway (BEER Run) experimental paradigm reveals *Drosophila* travel more direct paths to receive 50% ethanol. (**A**) Schematic diagram of BEER Run operant administration assay. (**B**) Side-view schematic of one-of-six lanes within BEER Run apparatus depicting the five main stages of this operant administration assay: (*i*) acclimation within the start chamber, (*ii*) time out in the runway, (*iii*) volatized-ethanol or air administration in the end chamber, (*iv*) time out in the runway, and (*v*) fifty-minute inner-interval period in the start chamber. (**C**) Previously mentioned stages represent one trial of operant administration within the full experimental paradigm across two days that includes six trials. Tracking data was not collected during start chamber acclimation, the time out in runway directly following ethanol exposure, or the fifty-minute inner-interval periods within trials. (**D**) Example automated tacking results of fly coordinate position during time out in the runway for trials 1-6 recorded at 15 frames per second. (**E**) Average angular velocity counts were recorded for six spaced operant administration trials over two-days at varying concentrations of ethanol (*i*. 30%; *ii*. 40%; *iii*. 50%; *iv*. 80; *v*. 85%; *vi*. 90%). Average angular velocity during the operant task was collected for each fly in air and ethanol receiving groups and plotted across six trials standardized to trial one behavior. Square-points indicate group mean +/- SE. Repeated ANOVA with planned contrasts for each concentration 30%, 40%, 50%, 80%, 85%, 90% indicate that (*i*) 30% ethanol (air n = 5, ethanol n = 6) and (*ii*) 40% ethanol (air n =10, ethanol n = 12) groups show significant differences on day two (30% F(2,18) = 4.08, p = 0.03; air v. ethanol T3, T4 p=0.011, 0.047; 40% F(2,40) = 4.73, p = 0.01, trials T1,T3 p=0.031, T4,T5 p=0.001, T4,T6 p=0.003) If Mauchly’s test indicated that the assumption of sphericity had been violated, the Greenhouse-Geiser correction was applied to the data. All posthoc analysis was performed with Bonferroni corrections. (*iii*) 50% ethanol group (air n = 23, ethanol n = 19) show significant differences on both days (F(2,80) = 12.06, p = 0.00; air v. ethanol T2,T5,T6 p=0.029, 0.036, 0.022, ethanol group T1,T2 p=0.007, T4,T5 p=0.048, T4,T6 p=0.018). (*iv*) 80% ethanol group (air n = 23, ethanol n = 30) did not show significance on either day (F(1.77,90.24) = 0.18, p = 0.81). (v) 85% ethanol (air n = 22, ethanol n = 22) and (vi) 90% ethanol (air n = 12, ethanol n = 9) groups shows significant differences predominantly on day 1 (85% F(2,84) = 7.95, p = 0.00; trials T1,T3 p=0.004, T4,T6 p=0.026; 90% F(2,38) = 5.73, p = 0.01; air v. ethanol day 1 p=0.048, T1,T2,T6 p=0.012, 0.005, 0.015, air group T1,T2 p=0.004, ethanol group T2,T3 p=0.028). See also Figure S1.(**F**) A heatmap of behavioral dynamics comparing the 50% ethanol group to the air group (air n = 23, ethanol n = 19) for all trials. Rows correspond to a behavior (angular velocity, velocity, latency, distance, time) and columns correspond to trials 1-6. Each grid corresponds to a behavior index calculated by first scaling and centering (mean = 0, sd = 1) behavior data for each fly within the ethanol and air receiving groups, and then calculating a behavior index for each behavior across trials: 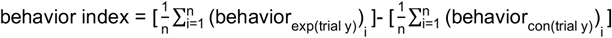. Color indicates how much more (red) or less (blue) the behavior occurred in the ethanol group compared to the air group. See also Figure S2.(**G**) Significant trials identified per behavior. 50% ethanol group shows significant differences in (Eiii, F, Gi) angular velocity (reported above) and (F, Gii) velocity (MANOVA, Wilks’ Lambda day*trials*group: V = 0.86, F(2,39) = 3.16, p =0.05, ω^2^ = 0.14; air v. ethanol T2,T6 p=0.005, 0.011, air group T1,T2 p=0.004, T1,T3 p=0.000, T3,T6 p=0.05, ethanol group T1,T2 p=0.000, T1,T3 p=0.002, T4,T6 p=0.004). See also Figure S2.

In order to pursue these questions, we defined parameters within BEER Run such that the consequence of a single-fly traversing the runway and entering the end chamber (runway task) was a defined dose of volatilized ethanol (Figure 1B). Each fly was presented with this choice three times per day for two consecutive days (Figure 1C). The ethanol receiving flies were compared to age-matched control flies that received air in the end chamber upon runway task completion. To quantify fly behavior, the BEER Run apparatus is paired with robust, high resolution tracking software that tracks fly coordinate position (x,y) at 15fps (Figure 1D). BEER Run and the tracking software are integrated and controlled through an easy-to-use graphical user interface (GUI) (Additional Supplementary Item 1). This tracking data in combination with Ctrax computer vision [10] and JAABA machine learning [11] open source software allowed us to consider 25 different behaviors and behavioral features during the runway task and ethanol administration.

### Dose responsivity of ethanol as a reinforcer

Individual flies were exposed to a fixed concentration of volatized ethanol (30%, 40%, 50%, 80%, 85% or 90%) or humidified air for 10min at 30 cm^3^/min upon end chamber entry for all 6 trials. Coordinate position measurements at 15fps were converted to operant task metrics including angular velocity, velocity, distance, latency, and time to traverse runway (Figures 1E-1G, S1 and S2). Angular velocity (aka turning velocity) during the operant task showed differences across trials at 3 ethanol concentrations when compared to flies receiving air in the end chamber. These concentrations include a range of low (30% ethanol: air n = 5, ethanol n = 6), moderate (50% ethanol: air n =23, ethanol n = 23) and high (90%: air n = 12, ethanol n = 9) ethanol concentrations (Repeated Measures ANOVA with Posthoc Bonferroni corrections: <u>30%</u>, F(2,18) = 4.08, p = 0.03; air v. ethanol T3, T4 p=0.011, 0.047, air group T2,T3 p=0.039, T3,T6 p=0.049, ethanol group T1,T4 p=0.031; <u>50%,</u> F(2,80) = 12.06, p = 0.00; air v. ethanol T2, T5, T6 p=0.029, 0.036, 0.022, ethanol group T1,T2 p=0.007, T4,T5 p=0.048, T4,T6 p=0.018; <u>90%,</u> F(5,95) = 5.069 p=0.000; air v. ethanol T1, T2, T6 p=0.012, 0.005, 0.015, air group T1-T2 p=0.018, T1-T6 p=0.040) (Figures 1E-1Gi). Only in the 50% ethanol group did angular velocity show significant differences between groups regarding all of day 1 (p=0.033) and day 2 (p=0.007) and decrease throughout the course of each day, indicating more directed paths to ethanol administration. Comparing velocity on subsequent trials to T1 demonstrated that administration of 50% ethanol decreased operant task velocities on trial 2 and trial 6 compared to air controls (MANOVA F(2,39) = 3.16, p =0.05; Wilk’s V = 0.86, ω^2^ = 0.14; T2, T6 p=0.005, 0.011) (Figures 1F, 1Gii, S1Aiii, and S2C, S2E, Additional Supplemental Item 2).

For 50% ethanol, latency to leave the end chamber and time to complete runway task were not significantly altered by ethanol exposure (Figure 1F and S1). However, high concentrations (85%, 90%) of ethanol increased latency on day 1 (<u>85%:</u> F(1.33,55.76) = 3.91, p = 0.04,Greenhouse Geisser correction ε=0.66; day 1 p=0.010, T2,T3,T5 p=0.006, 0.053, 0.024; <u>90%:</u> F(1.11,21.17) = 4.73, p = 0.04; air v. ethanol p=0.025, day 1 p=0.026, T2 p=0.027) and increased time taken to complete the task on day 2 (90%: F(2,38) = 3.77, p = 0.03; air v. ethanol p=0.033, day 2 p=0.046). Increased latencies on day 1 and increased times on day 2 cumulatively indicate learned aversion for ethanol, suggesting that high concentrations may confound the ability to detect appetitive behavior in this two-day paradigm.

### Behaviors demonstrated during ethanol exposure

To examine behaviors demonstrated during 50% ethanol exposure, coordinate position measurements at 15fps were converted to velocity (Figure 2A-F). Further we ran experiment video files through computer vision (Ctrax) [10] and machine learning software (JAABA) [11] to quantify a series of custom behaviors (pausing, pacing, thrashing) in a randomized subset of the data (50% ethanol, n =12; air, n = 12). Power analysis for k=2 groups, a significance level of 0.05 and a power of 0.8 indicated that an n=8 would be sufficient to determine differences between groups. 50% ethanol exposure significantly decreased velocity on the first exposures of each day (F(1.58,63.30) = 28.18, p = 0.00, Greenhouse Geisser Correction ε=0.79; T1, T4 p=0.000, 0.003) (Figures 2A and 2D). However, consecutive exposure to ethanol resulted increased velocity on the last trial of day 2 (T6, p=0.027).

**Figure 2.**
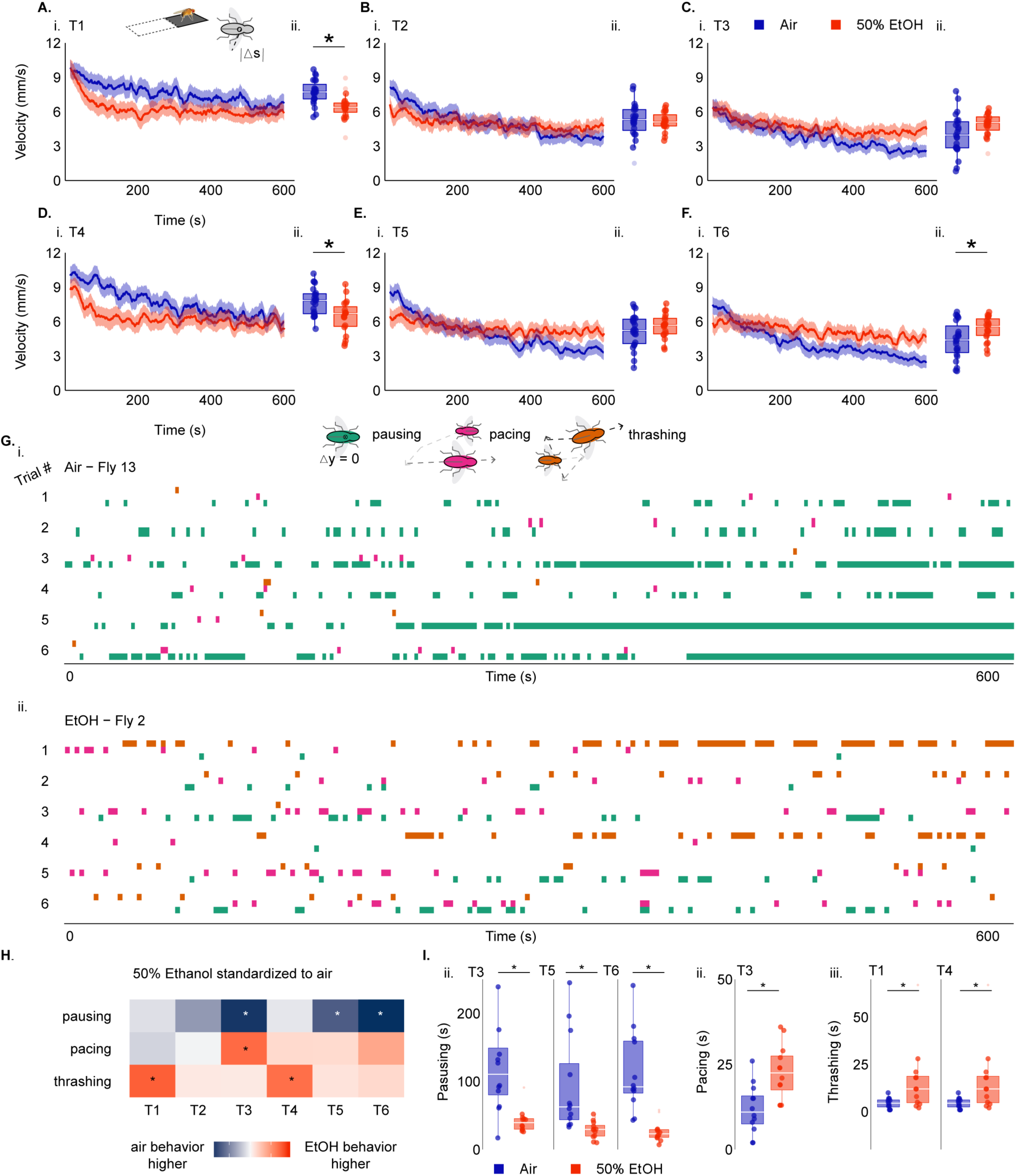
*Drosophila* behaviors during initial ethanol experiences. (**A-F**) Velocity counts for each trial were binned over 15 second periods, averages across biological replicates (air n = 23, ethanol n =19), and plotted against time. Lines depict mean +/- SE. (**Aii, Bii, Cii, Dii, Eii, Fii**) Average group velocity across 600s of exposure is recorded as boxplots. Repeated measures ANOVA with planned contrasts and posthoc Bonferroni corrections indicate a significant interaction between ethanol administration velocity and trial number (F(1.58,63.30) = 28.18, p = 0.00, Greenhouse Geisser Correction ε=0.79; air v. ethanol T1, T4, T6 p=0.000, 0.003, 0.027). Ctrax version 0.3.1 computer vision software paired with “fixerrors-0.2.23” protocol extension were used to collect more detailed trajectory information for 24 flies (50% ethanol, n =12; air, n = 12). These videos and trajectory data were paired with three behavioral classifiers (pausing, pacing, thrashing) previously trained in this laboratory using supervised machine learning, JAABA. (**G**) Representative behavioral classifier ethograms that depict pausing (green), pacing (pink) and thrashing (orange) occurrences for six-trials over 600s of exposure to air (*i*) and 50% ethanol (*ii*). (**H**) A heatmap of behavioral dynamics comparing behavior during 50% ethanol exposure to air exposure (air n = 12, ethanol n = 12) for all trials. Rows correspond to a behavior (pausing, pacing, thrashing) and columns correspond to trials 1-6. Each grid corresponds to a behavior index calculated as previously stated in Fig.1. Color indicates how much more (red) or less (blue) the behavior occurred in the ethanol group compared to the air group. See also Figure S3. (**I**) Significant administration trials identified per behavior. Repeated measures ANOVA with planned contrasts and posthoc Bonferroni corrections indicate that the 50% ethanol group shows significant differences in administration behavior regarding (H, Ii) pausing (F(2,44) = 14.09, p = 0.00; air v. ethanol p=0.001, day 1, 2 p=0.004, 0.001, T3, T5, T6 p=0.001, 0.006, 0.000, air group T1,T2 p=0.004,T1,T3 p=0.000, T2,T3 p=0.001, T4,T5 p=0.001, T4,T6 p=0.000, T5,T6 p=0.041), (H,Iii) pacing (F(2,44) = 9.56, p = 0.00; air v. ethanol T3 p=0.002, air group T1,T4 p=0.044, ethanol group T1,T3 p=0.015), and (H,Iiii) thrashing (F(1.17,25.74) = 5.77, p = 0.02; air v. ethanol p=0.023, day 1,2 p=0.033, 0.048, T1, T4 p=0.031, 0.020, ethanol group T1,T2 p=0.005, T1,T3 p=0.016, T4,T5 p=0.002, T4,T6 p=0.002). See also Figure S3.

Behaviors obtained through machine learning, pausing, pacing, and thrashing distinctly changed due to repeated ethanol exposure (Figure 2G-2I and S3). During the later trials of each day the 50% ethanol receiving flies showed significantly less pausing (F(2,44) = 14.09, p = 0.00; day 1, 2 p= 0.004, 0.001, T3,T5,T6 p=0.001, 0.006, 0.000) (Figure 2G, 2H, 2Ii and S3B). Pacing behavior increased over the course of each day, and was significantly increased on the last trial of day one (F(2,44) = 9.56, p = 0.00; T3 p=0.015) (Figure 2G, 2H, 2Iii and S3C). During the first trial of each day, ethanol significantly increased thrashing behavior (F(1.17,25.74) = 5.77, p = 0.02, Greenhouse Geisser Correction ε=0.67; day 1,2 p = 0.033, 0.048, T1,T4 p=0.031, 0.020) (Figure 2G, 2H, 2Iiii and S3D).

Analysis of ethanol body content in these flies indicate that immediately following 10min of 50% ethanol exposure, flies contain 42.81 +/- 9.69 mM/fly ethanol, and following 50min of rest during the inter-interval trial internal ethanol concentration is significantly reduced to 6.70 +/- 4.22 mM/fly (Independent Group t-test t(16) = 3.976, p = 0.002) (Figure S3E-G). Importantly, the ethanol body content following 50min rest is not significantly different from air control levels (Independent Group t-test t(16) = - 1.214, p = 0.242), indicating that ethanol receiving flies have time to fully metabolize ethanol in between operant tasks (Figure S3F).

### Behaviors demonstrated during ethanol seeking

To better understand the complex behavioral dynamics that result in the development of seeking behavior despite the aversive properties of ethanol, Ctrax [10] and JAABA [11] were applied to videos of flies during the runway operant task (Figure 3A). The 15 per-frame features and 5 behavioral classifiers used to quantify fly behavior (50% ethanol, n =12; air, n = 12) revealed the dynamic nature of behavioral features across trials (Figures 3B-D, S4, S5 and Additional Supplementary Item 3). Upon first entry into BEER run flies explore the full apparatus at the fastest times exhibited within this paradigm (23.48 +/- 1.24 seconds). Thus, we split operant task into 2 phases: the time spent in the start chamber before initiating the task (start chamber behavior - Figure 3C), and the last 25 seconds of the trial (Figure 3B). Analysis of the time in the start chamber revealed that ethanol group flies generally faced walls more before initiating the task (T2,T3,T6 p=0.030, 0.055, 0.010) (Figure S5A). In addition to significant differences seen in *angle2wall* (T1,T6 p=0.015, 0.007) and *dphi* (T3,T4 p=0.001, 0.055) during the entire operant trial (Figure S5B), the last 25 seconds of the task included significant differences in speed and turning related features: *angle2wall* (T1, T6 p=0.016, 0.000), *dphi* (T3 p=0.015), *signdtheta* (T1 p=0.019), and *theta* (T2 p=0.013) (Figure 3B, S5B, S5C).

**Figure 3.**
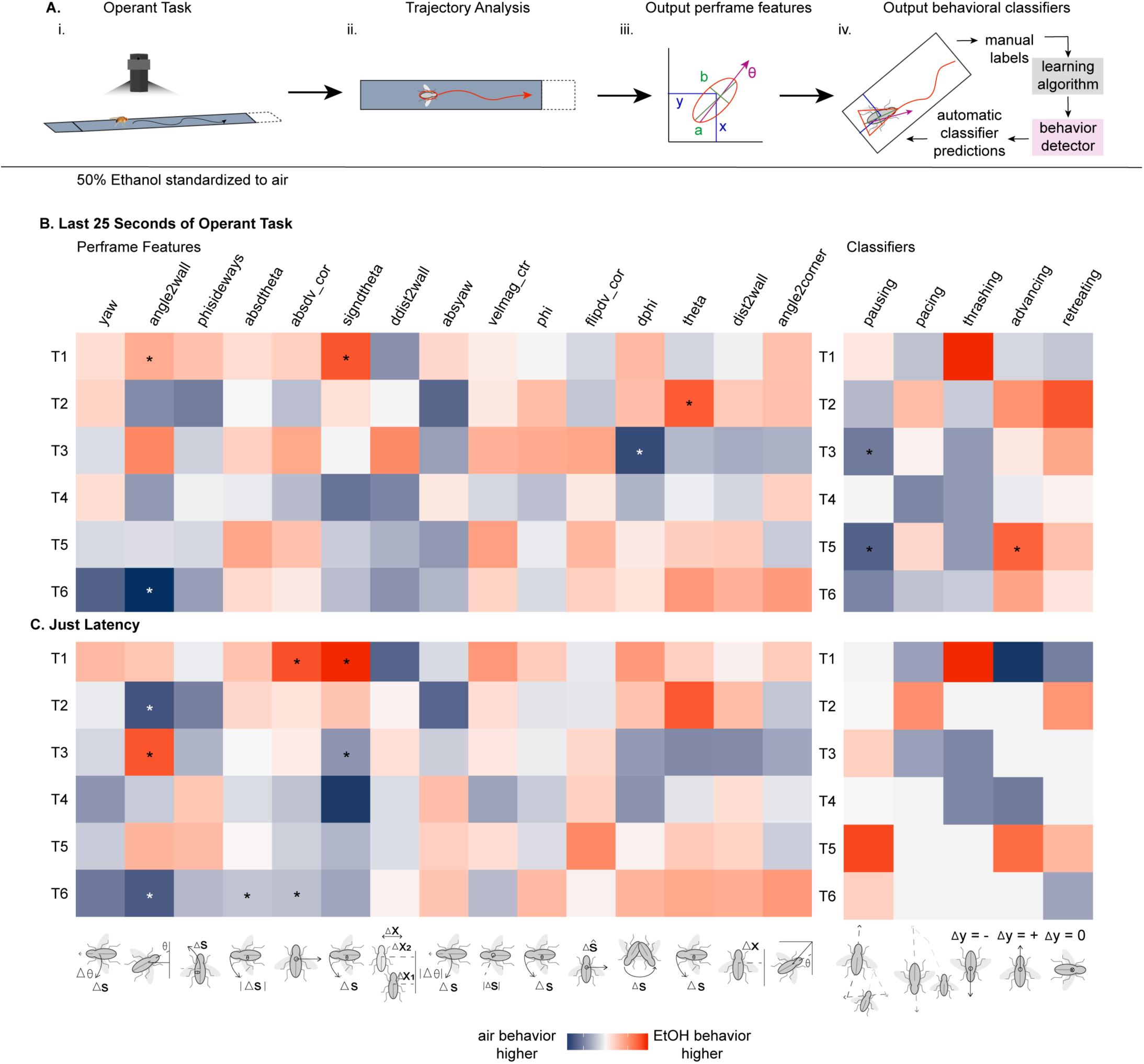
Behavioral features and classifiers associated with progressive ethanol administration. (**A**) Behavioral analysis pipeline including the (*i*) BEER run operant task, (*ii*) detailed trajectory analysis using Ctrax version 0.3.1 software, (*iii*) perframe feature behavioral output that is used to inform machine learning (*iv*) behavioral classification. This pipeline was used to collect behavioral counts for 24 flies (50% ethanol, n =12; air, n = 12). (**B-C**) Heatmaps for the last 25 seconds of the task (B) and the period before task initiation (C) were constructed to compare behavioral dynamics in 50% ethanol group to the air group. Heatamps include behavioral perframe features: angle2corner, dist2wall, theta, dphi, flipdv_cor, phi, velmag_ctr, absyaw, ddist2wall, signdtheta, absdv_cor, absdtheta, phisideways, angle2wall, yaw (see also Figure S4), and behavioral classifiers: pausing, pacing, thrashing, advancing, and retreating. Each grid corresponds to a behavior index calculated as previously stated in Fig.1. Color indicates how much more (red) or less (blue) the behavior occurred in the ethanol group compared to the air group. (**B**) Repeated measures ANOVA with planned contrasts and posthoc Bonferroni corrections indicate significant differences during the last 25 seconds of the operant task included significant differences in angle2wall (F(2,44) = 4.54, p = 0.02; air v. ethanol day 2 p=0.001, T1, T6 p=0.016,0.000, air group T1,T4 p=0.005, T3,T6 p=0.000, T5,T6 p=0.005), signdtheta (F(1,22) = 6.52, p = 0.02; air v. ethanol T1 p=0.019, air group T1,T4 p=0.021), dphi (F(2,44) = 4.00, p = 0.03; air v. ethanol T3 p=0.015, air group T4,T5 p=0.044, ethanol group T3,T6 p=0.014), theta (F(2,44) = 3.36, p = 0.04; air v. ethanol T2 p=0.013, air group T1,T2 p=0.005, T2,T3 p=0.040), pausing (MANOVA, Wilks’ Lambda trials*group: V = 0.97, F(2,21) = 0.25, p =0.023, ω^2^ = 0.30; air v. ethanol p=0.008, day 2 p=0.01, T3, T5 p=0.05, 0.03, air group T1,T2 p=0.003, T1,T3 p=0.000, T4,T5 p=0.000, T4,T6 p=0.000, ethanol group T1,T2 p=0.036, T1,T3 p=0.001, T4,T5 p=0.042, T4,T6 p=0.023), and advancing (F(2,44) = 3.22, p = 0.05; air v. ethanol T5 p=0.032). (**C**) The latency period before initiating the trial included significant differences in angle2wall (F(2,44) = 6.18, p = 0.00; air v. ethanol T2,T3,T6 p=0.030, =0.055, 0.010, air group T1,T2 p=0.017, T2,T3 p=0.003), absdtheta (F(1,22) = 4.36, p = 0.05; air v. ethanol T6 p=0.032), and signdtheta (F(1.59,34.96) = 4.11, p = 0.03; air v. ethanol day 2 p=0.024, T1,T3 p=0.019, 0.038, air group T1,T4 p=0.035). Significant differences were not observed for other behavioral features and behavioral classifiers. See also Figure S5. For per-frame feature descriptions see Additional Supplement Table S3.

Based on speed and turning behaviors that were most dynamic and significant across trials (*angle2wall, signdtheta, phi, theta, absdv_cor, absdtheta*) (Figure S5A-C, S5A, S4) and previous findings suggesting stagnant wall behavior is associated with avoidance [12], we hypothesized that behavioral features that incorporate turning, speed and position in the arena would be important for quantifying motivated response. Turning related behavioral features revealed that flies from the both the ethanol and air groups did not have *signdtheta* values significantly different from 0 for all trials following initial exposure (F(1,22) = 6.96, p = 0.02, T1 p=0.023), indicating that flies generally turned left and right equally (Figure S4 and S5A). No significant differences were seen between groups in the following speed related behavioral features *phisideways* (F(2,44) = 1.70, p = 0.19), *yaw* (F(2,44) = 0.32, p = 0.727), *absdtheta* (F(2,44) = 0.31, p = 0.74), *signdtheta* (F(2,44) = 0.28, p = 0.18), *flipdvcor* (F(2,44) = 1.78, p = 0.76), *absdv_cor* (F(2,44) = 0.96, p = 0.39), *phi* (F(2,44) = 0.34, p = 0.71), *absyaw* (F(2,44) = 1.11, p = 0.34), and *velmag* (F(2,44) = 0.15, p = 0.86) (Figures 3B and S4). Similarly, position in arena features including *dist2wall* (F(2,44) = 1.05, p = 0.36), and *angle2corner* (F(1.50,32.90) = 2.46, p = 0.11, Greenhouse-Geisser Correction ε=0.75) show no significant differences (Figure 3B and S4).

To further investigate which behaviors were learned responses to the appetitive and aversive properties of ethanol, we investigated how ethanol affected pausing, advancing, retreating, pacing and thrashing (Figures 3B, 3C and S4). Pausing, pacing and thrashing were chosen due to human observed behaviors and classified using JAABA. Advancing and retreating behaviors were classified to determine if flies exhibited approach-avoidance conflict [13] seen in other runway models [2]. Interestingly, on day 2 the ethanol group flies showed significantly decreased pausing (air v. ethanol p=0.008, day 2 p=0.01, T3,T5 p=0.05, 0.03) and increased advancing (T5 p=0.032) during the last 25 seconds of the operant task (Figure 3B), supporting appetitive goal-seeking behavior. No significant differences were seen between groups for retreating (F(2,44) = 0.50, p = 0.61), pacing (F(2,44) = 0.06, p = 0.94), and thrashing (F(1.30,28.69) = 3.01, p = 0.08, Greenhouse Geiser Correction ε=0.65) behaviors (Figure S5A).

### Combinations of behavioral features that define motivation for a complex stimulus

Our data thus far suggests flies demonstrate behavioral complexity as valence for ethanol shifts from aversive to appetitive. Observation of how behaviors change over time suggests that behaviors occur in combination (Figure 3), and clusters of behaviors may be more useful for defining a motivated response. To visualize how behaviors cluster together, we summed occurrences of behavior over the six trials to obtain a single metric for each behavior and constructed behavior matrices for air (Figure 4A) and ethanol (Figure 4Aii) treated flies. We observed that behavioral contributions differed in the air and ethanol conditions with *angular velocity and absyaw; velocity and absdtheta; latency, angle2wall, advancing*, and *pausing* being positively correlated with each other and strong negative correlations between *pausing* and *absdtheta*; *theta* and *angle2wall* in the ethanol condition (Figure 4A).

**Figure 4.**
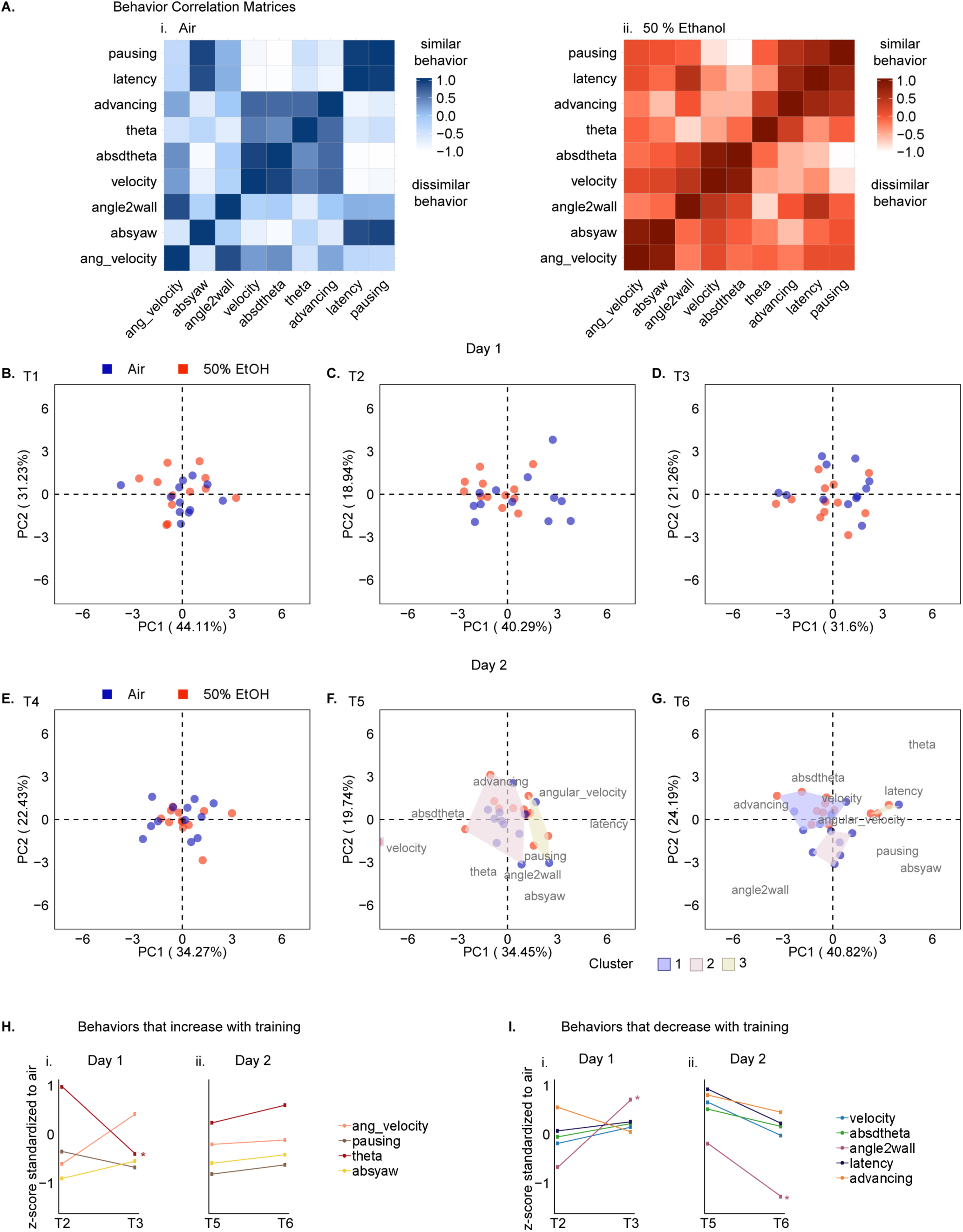
Motivation defined by a subset of behaviors highlights the switch from development to reinforcement learning on day two. (**A**) Behavior correlation matrices display the significance level between average z-scored behaviors collected across six trials for 24 flies (50% ethanol, n =12; air, n = 12). (ii) 50% ethanol correlation matrix was arranged using hierarchical clustering with linkage criterion = Ward’s, and the (i) air correlation matrix was arranged to compliment. (**B-G**) PCA plots were constructed for six-trials, where each fly was plotted as a function of 9 behaviors (pausing, latency, advancing, theta, absdtheta, angle2wall, absyaw, velocity and angular velocity). Behavior matrices for PCA plots were constructed from scaled and centered (mean = 0, sd = 1) behavioral data for each fly and each trial. From these data, euclidean distance matrices were constructed for each of the 6 trials, where observations = 24 flies, and features = 9 behaviors. Each trial’s distance matrix was projected onto its first 2 principal components. Color indicates ethanol receiving (red) and air receiving (blue) flies. (**B-E**) PCA plots for trials 1-4 do not suggest significant differences between ethanol fly behavior and air fly behavior. (**F**,**G**) Agglomerative hierarchical clustering was performed on trial 5 (**F**) and trial 6 (**G**) with linkage criterion = Ward’s. The sum of the within-cluster inertia was calculated for each partition. Clusters were defined from the partition with the higher relative loss of inertia (i(clusters n+1)/i(cluster n)). (**F**) trial 5 clustering indicates 50% of ethanol flies (6 of 12) cluster together driven primarily by behavioral contributions such as velocity and advancing. (**G**) trial 6 clustering indicates 75% of ethanol flies (9 of 12) cluster together in an upper quadrant cluster driven by behavioral contributions such as advancing and theta [see S4]. The same 24 flies (50% ethanol, n =12; air, n = 12) were used to plot (**H**) behaviors that increase with training and (I) behaviors that decrease with training. Each point reflects a behavior index that was calculated by first obtaining z-score (mean = 0, sd = 1) behavior data for each fly within the ethanol and air receiving groups, and then standardizing ethanol z-scores to the average of air controls for each behavior. Positive slope indicates that the behavior increased in the experimental group that day, while negative slope indicated that the behavior decreased in the experimental group that day. Paired T-Tests were performed on every behavior comparing the last two trials of day one (T2-T3) and the last two trials of day two (T5-T6). Tests reveal significant differences in (Hi) theta (T2,T3 t(11) = 3.13, p = 0.01), (Ii) angle2wall (T2,T3 t(11) = -4.12, p = 0.00), (Iii) angle2wall (T5,T6 t(11) = 3.46, p = 0.01). For per-frame feature descriptions see Additional Supplement Table S3. For more information on PCA see Additional Supplement Item 4.

To visualize the impact of multiple behaviors during the runway task, principal component analysis (PCA) was used to reduce animal behavior exhibited during trials onto 2 principal components, and cluster analysis was used to identify groups of similarly behaving animals. Nine of the most dynamic behaviors that highlighted significant differences between air and ethanol groups were selected for PCA analysis including *angular velocity, velocity, advancing, pausing, theta, absdtheta, angle2wall*, and *absyaw* (Figures 4, S4, and S5). There was a large overlap in fly behavior for ethanol group flies and air group flies in trials 1 (Figure 4B), 2 (Figure 4C) and 3 (Figure 4D). This trend was anticipated because during the first day of exposures ethanol receiving flies are learning to associate the end chamber with ethanol and the consequences of ethanol intoxication. By trial 4 the ethanol flies begin to group tighter together (Figure 4E). On trial 5, 50% of ethanol flies cluster together driven by *velocity* and *advancing* which contribute to PC1 by -0.791 and PC2 by 0.362 respectively (Figure 4F, Additional Supplemental Item 4). The grouping is most significant on trial 6, where 75% of ethanol flies cluster together in the upper quadrant cluster driven predominantly by *advancing* and *theta*, which contribute to PC1 by -0.512 and PC2 by 0.587 respectively (Figure 4G, Additional Supplemental Item 4).

This data suggests a shift in behavior between Trials 5 and 6 from aversive to appetitive behavior. To highlight the behavioral dynamics driving the formation of these clusters over trials, we compared the patterns of behaviors between Trials 5-6 to Trials 2-3 (Figures 4H and 4I). Behaviors that tended toward an increase in between Trials 5 and 6, including *angular velocity, pausing, theta*, and *absyaw* did not show a similar pattern between Trials 2 and 3. Similarly, behaviors that trended toward a decrease between Trials 5 and 6 including *velocity, absdtheta, latency, advancing and angle2wall* (paired t-test: angle2wall T5,T6 t(11) = 3.46, p = 0.01), did not show a similar pattern between Trials 2 and 3 (Figure 4I).

## Discussion

Flies exhibit preference for an odor associated with moderate dose ethanol intoxication one to seven days after pairing [14, 15]. However, this Pavlovian assay lacks the tracking resolution to quantify behavior during ethanol administration. Operant runway models are commonly used as a tool for studying goal-seeking motivated behavior [2, 9, 16]. An unfortunate disadvantage of this model is that it can be difficult to assess how much effort the animal is expending to achieve the reward [2, 9]. Our assay overcomes this disadvantage by using high content tracking which adds a rich repertoire of behavioral metrics that can be associated with different aspects of the task in this apparatus. Moreover, the high-content behavioral data allowed us to identify the subtle behavioral dynamics contributing to the shift from ethanol aversion to preference.

Our runway-based operant administration data here reveal that flies willingly self-administer ethanol following initial exposure, despite finding the ethanol aversive. Interestingly, the ethanol dose that induced the most striking behavioral responses in the runway task is similar to the one that induced the highest appetitive response in a Pavlovian odor-ethanol association memory task [15]. We found that within each day, initial ethanol administration induced early bouts of thrashing behavior, followed by decreasing velocity and increased levels of pausing, cumulatively supporting aversion. Remarkably however, even though the ethanol exposure appeared aversive, flies readily self-administered 50% ethanol within 5 minutes of being presented with the choice. However, within later trials, flies pause significantly less while receiving ethanol, and while velocity remains steadily above control behavior, these flies begin pacing in the absence of thrashing. Together these behavioral data are suggestive of formation of behavioral ethanol tolerance with consecutive ethanol exposures.

We postulate this tolerance alters the perceived valence of the ethanol stimulus. Behavior in the runway suggests that ethanol was initially aversive, resulting in slower velocities and increased turning on day 1, followed by a switch in behavior on day 2 where flies travel more goal-directed paths to the end chamber. This transition from aversive to appetitive response appears to shift between Trial 5 and Trial 6, characterized by a combination of 9 behaviors that comprise the position, direction and movement of the animals.

Behavior during our runway task thus supports the biphasic actions of ethanol intoxication, which include initial ‘euphoria’ associated with reward induced by volatized ethanol administration, followed by negative ‘dysphoria’ as intoxication declines [17]. Presence of the biphasic action of ethanol coupled with the development of opposing behaviors during the runway task (e.g. decreased velocity and increased latency but decreased angular velocity) are supported by the opponent process theory, which states that self-administered drugs should have positive (euphoric) and negative (dysphoric) properties that can lead to classic “approach-avoidance” conflict [13, 18]. We speculate that similar conflict is represented in *Drosophila* by increased latency and decreased velocity corresponding with a more goal-directed path to the end chamber in our data. The BEER Run provides an ideal platform to test how the aversive consequences of drugs of abuse contribute to the drug seeking through reinforcement, thus contributing to a mechanistic understanding of the Opponent-Process Theory.

In *Drosophila* learned responses to aversive and appetitive stimuli are inextricably linked through valence modules in the mushroom body [19-21]. Moreover, aversive experiences can influence perception of rewarding stimuli in *Drosophila*. This can happen whether different aversive and appetitive stimuli are presented simultaneously [22], whether a single stimulus has both aversive and appetitive properties [14, 15], or whether a previous aversive experience alters internal state in the brain, thus affecting future appetitive experiences [23]. The BEER run provides an opportunity to investigate how these valences are integrated and reflect the appropriate output response.

Our data also suggest that *Drosophila* may be an appropriate model to investigate the mechanisms underlying incentive salience. A prominent theory in the field of addiction research is that cues that are associated with drugs can acquire features that mimic those produced by the drugs themselves and thereby serve to increase the effectiveness to trigger drug seeking [24, 25]. The cues thus gain incentive salience, which confers a desire or ‘want’ attribute. Alcohol-predictive cues can accrue incentive salience in rodent models [25]. *Drosophila* initiate seeking when exposed to a cue previously associated with alcohol and will also walk over a 120V electric grid to attain that cue [14]. We suggest the runway environment accrues incentive salience with consecutive trials since the flies express more goal-directed behavior despite demonstrating initial aversion to the volatilized ethanol. Whether flies increase responsivity to a discrete, localizable conditioned stimulus that initiates seeking can now be tested with this apparatus.

In summary, the BEER Run apparatus combined with live and offline tracking effectively provide a set of behavioral features associated with motivated response in *Drosophila*. Our high content behavior analysis demonstrates subtle behavioral changes associated with a shift in valence for ethanol from aversive to appetitive. Moreover, this apparatus can easily be adjusted for delivery of other substances, whether they be intoxicating drugs or natural appetitive and aversive substances. Understanding the combination of subtle behavioral features associated with operant memory in individual animals is a critical step forward for understanding the neural mechanisms underlying acquisition, consolidation and expression of these memories.

## Materials and Methods

### Experimental Model and Subject Details

Canton-S (CS) wild-type flies were reared at 25°C in humidity-controlled incubators under 14:10 Light:Dark (L:D) cycles. Flies were raised in 9.5cm (height) x 2.5 cm (diameter) polypropylene vials on standard cornmeal agar food media with anti-fungal agent. Male flies were collected 1-2 days post eclosion under light, humidified CO_2_ anesthesia and experiments initiated when flies were 3-5 days old. Flies were transferred via mouth-pipette prior to behavior experiments and sacrificed following each experiment.

### Method Details

#### BEER Run Hardware

The BEER Run apparatus is a custom-built six individual lane behavioral chamber. Each lane is divided in three sub-compartments referred to as the start chamber (13mm x 6mm), runway (10cm x 6mm) and end chamber (18mm x 6mm), which can be conditionally and temporally segregated by a gating system mechanically controlled by a metal gear servo (HS_5056MG, Hitec, Hazard Way Suite D, Sand Diego, CA, USA) (Figure 1A and Additional Supplementary Item 1). All mechanical gating systems were connected to the BEER run assay operating system (developed using ROS by Peter Polidoro, Senior Engineer, Janelia Research Campus). Assembled lanes had approximately 3 mm of headspace to prevent flight and promote free walking.

Each lane is made of virgin PTFE material (machined by Whiteglove Inc.) with a transparent top made of clear acrylic sheet laser cut to size. The odors, air and/or ethanol were introduced to each runway via small openings in the runway and were controlled by the three separate odor delivery system called olfactometers. Each olfactometer consist of one control panel that controlled two IQ+Flow digital mass flow meters (IQF-200C-AAS-00-V-S, 10psi(g)/0psi(g), Bronkhorst, Ruurlo, Netherlands), and two solenoid pinch valve assembly (InLine 5×2-way Normally close, NResearch Inc, West Cadwell, NJ) connected to five 40ml glass sample vials (27184, Millipore Sigma, Darmstadt, Germany). The complete three olfactometer assemblies were connected to the BEER run assay operating system [26]. Vapor flow was directed from vials through apparatus end chamber ports with 1/8” diameter PTFE tubing (PTFE Masterflex Tubing, 1/8” OD, Coleparmert, IL, USA). All lanes were individually connected to an olfactometer for air and volatized ethanol delivery; and an active vacuum with equal flow rate as the input flow to facilitate flow within the end chamber. An additional flowmeter was added at the entry point to the runway to monitor the input flow in order to regulate the 2^nd^ flowmeter that was connected to the vacuum lines (AALBORG, Orangeburg, NY, USA)

The fully assembled BEER Run apparatus consists of six custom built lanes, placed in an aluminum housing with clear acrylic sheet covering each lane. An additional narrow clear acrylic sheet provides a port to introduce flies into the runway. The aluminum arena holder (Custom made aluminum frame, JRC, Ashburn, VA, USA) is held by four posts providing stability and space for IR back light to illuminate the runway without any obstruction. A Basler Firewire Mono camera equipped with 25MM lens with IR filter (Edmund Optics Inc. East Bloucenter Pike, Barrington, NJ, USA) provides a clear and complete view of the arena. Each lane is equipped with two metal gear micro servos to be activated and trap the flies depending on the position of fly.

Odor ports in the runway near the start and end chamber gates are engineered to deliver humidified air to ensure the fly does not become dehydrated during extended protocols. Alternatively, they can be used to deliver an odor cue that becomes associated with the runway (Figures 1A and 1B). This allows simultaneous measurement of associative and operant memory or may be used to measure extinction or reinstatement to a reward-associated cue. Hardware is described in detail in Additional Supplemental Item 1.

#### BEER Run Behavioral Experiments

Operant Ethanol Administration Assay: For behavior experiments, flies were individually exposed to six operant administration trials over two days (Figure 1C). The assay was performed with lights on and temperature and humidity controlled at 25°C and 55% respectively. To start the assay one fly is placed in each start chamber, mechanically gated and segregated from the rest of the lane, for a 5 min acclimation period. Post acclimation, the training trial was initiated by start chamber and end chamber gates opening allowing full access to the lane. The fly then had a maximum of 5 min to traverse the runway and enter the end chamber, which triggered mechanical gate closure. This initiated 10 min of vaporized ethanol administration within the end chamber. Following administration, start chamber and end chamber gates open and a flashing blue LED (1 flash/sec for 10 sec at 20% capacity) along the rear wall of the start chamber encouraged the intoxicated fly to re-enter the start chamber, with a 5 min time cap. Entry into the start chamber triggered mechanical gate closure, initiating a 50 min inter-trial interval wait period. Each fly is exposed to this paradigm two more times on day one, individually housed overnight in food vials at 25°C in humidity-controlled incubators under 14:10 Light: Dark (L:D) cycles, and then exposed to three more trials on day two. Ports located in the start chamber and end chamber delivered humidified air when vaporized ethanol was not being administered in the end chamber to ensure that flies did not dehydrate during experiments.

We developed an easy-to-use GUI to adjust experimental settings (Additional Supplementary Item 1). While the experimental settings are flexible and can accommodate many different behavioral protocols the optimized protocol for this operant assay includes 6 trials over 2 days, a 5min time cap to traverse the runway and enter the end chamber associated with ethanol, a 10min vaporized ethanol exposure duration, and a 50min inner-trial interval (Figures 1A-C). Ethanol solutions were prepared fresh for each two-day experiment. For example, a 50% ethanol solution: 50mL Ethyl Alcohol 200 proof (Pharmaco by Greenfield Globe, Brookfield, Connecticut, USA) + 50mL ddH_2_O.

#### Ethanol Absorption and Metabolism

Flies were frozen in liquid nitrogen immediately following 10min of 50% ethanol exposure for absorption studies and 50min post-exposure for metabolism studies. Flies were homogenized in 0.5M glycine (pH 9, 200µl for 6 flies). 2-fold dilution was performed on 100µl of supernatant following centrifugation at 12,000rpm for 5min. Following a second round of centrifugation (12,000rpm for 5min) ethanol concentrations were measured in fly homogenate supernatant using NAD-ADH Reagent (N7160, Sigma-Aldrich, St. Louis, MO, USA). The reaction was placed in a Synergy HTX plate reader (BioTek, Winooski, VT,USA) and measured every 5 minutes for 30minutes at 340nm. To calculate the ethanol concentration in flies one male fly was estimated to be 0.75mg based on average mass recorded during experiments. Calculations involved extrapolating XmM ethanol content at 15min and a final calculation of 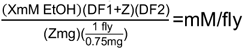, where DF is dilution factor and Z is total mass of flies within each sample.

#### Analysis of *Drosophila* Behavior

During experiments the Basler Firewire camera integrated with the BEER run assay operating system live-tracked fly position during pre-exposure time out in runway and ethanol administration in end chamber as (x, y) coordinates at 15 fps for all 6 lanes (Fig. 1c). Output files included a “.bag” video file and “tracking.txt” file for each trial. All coordinate position tracking data was processed in R studio using dplyr and reshape2 packages to determine latency to leave the start chamber (s) and pursue ethanol, time (s) taken to traverse the runway and enter the end chamber, as well as distance traveled (mm), velocity (mm/s), and angular velocity (deg/s) during both the runway task and ethanol administration.

#### Characterization of Detailed Trajectory Information

Trajectories of twenty-four flies (50% ethanol, n = 12; air, n = 12) were tracked using Ctrax version 0.3.1 computer vision software paired with “fixerrors-0.2.23” protocol extension to obtain more detailed trajectory information during the runway task and ethanol administration [10]. All BEER run “.bag” video files were converted to “.avi” files to maximize compatibility with Ctrax. In this method, an ellipse is fit to the body of each fly frame-by-frame through the use of background subtraction to classify pixels as a fly body or background. The Gaussian distribution determined through Expectation-Maximization for Gaussian mixture models is used to estimate an ellipse for the body of each fly, and orientation is estimated through a dynamic programming algorithm. Supervised tracking was used to correct ambiguous trajectory sequences. This pipeline was carried out in a blinded fashion, and behavioral features that measured individual fly behavior were selected for analysis: *absdv_cor, absdtheta, angle2corner_rect, angle2wall_rect, absyaw, ddist2wall_rect, dist2wall_rect, dphi, flipdv_cor, phi, phisideways, signdtheta, theta, velmag_ctr, yaw* (Additional Supplementary Item 3).

#### Characterization of Classified Behaviors

The interactive Janelia Automatic Animal Behavior Annotator (JAABA) machine learning tool [11] was used in order to obtain data on more complex behaviors: advancing, retreating, pausing, pacing and thrashing. To generate classifiers for each of these behaviors, a subset of videos (from air and ethanol receiving flies) was manually annotated with positive examples of frame-sequences in which a fly was performing the behavior of interest and negative examples of when a fly was not performing the behavior. A brief description of each behavior classified is listed below. Supervised behavioral annotations were processed by the JAABA machine learning algorithm’s interaction with trajectory data from Ctrax to produce behavioral classifiers for automatic annotation.

> Advancing – the fly, oriented toward the End Chamber, moves toward the End Chamber.
>
> Retreating – the fly, oriented toward the Start Chamber, moves toward the Start Chamber.
>
> Pausing – the fly has almost no translation or rotational body movement.
>
> Pacing – the fly has repetitive translational and rotational body movement while oriented toward an arena wall or arena gate. Thrashing – the fly performs large and rapid changes in rotation and/or translation for >1 second.

To quantitatively determine the generalization error of classifiers across the dataset, we used JAABA software’s “groundtruthing mode” to compare automatic behavioral classifiers to manually labeled data. If needed classifiers were rebuilt with additional supervised annotations until they performed with 100% accuracy during “groundtruthing mode”.

#### Analysis and Characterization of *Drosophila* Activity

All data from BEER run base tracking, Ctrax computer vision, and JAABA machine learning was processed in R studio using dplyr and reshape2 packages. Average behavior within each trial was calculated for each group and plotted with standard error. Most graphs include visual representation of individual fly data within groups.

Behavioral dynamics across trials was assessed using heatmap color gradient representations of behavioral differences in ethanol group v. air control group. Behavior differences were calculated by first scaling and centering (mean = 0, sd = 1) behavior data for each fly within the ethanol and air receiving groups, and then calculating a behavior index for each behavior across trials: 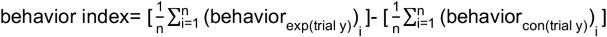

Principal component analysis and cluster analyses were conducted to visualize how fly behavior changed across trials, and to determine which behaviors were required for reinforcement learning and motivation for ethanol. The subset of 9 behaviors required for visualization of learning included *pausing, latency, advancing, theta, absdtheta, angle2wall, absyaw, velocity and angular velocity*. These behaviors were summarized by the average occurrence or summation of occurrences for each trial (fly identity is maintained). Behavior matrices were constructed to visualize each behavior across all 6 trials (fly identity is maintained). Each behavior matrix was scaled and centered (mean = 0, sd = 1). From these data, Euclidean distance matrices were constructed for each of the six trials, where observations = 24 flies and features = 9 behaviors. Each trial’s distance matrix was projected onto its first two principal components, where each fly was plotted based on the 9 behaviors it exhibited. Agglomerative hierarchical clustering was performed on trial 5 and trial 6 with linkage criterion = Ward’s. The sum of the within-cluster inertia was calculated for each partition. Clusters were defined from the partition with the higher relative loss of inertia (i(clusters n+1)/i(cluster n)).

#### Statistics and Data Visualization

Repeated Measures ANOVAs with planned contrasts were performed on all data to interpret significance between groups and across trials (Table S2). Mauchly’s test of sphericity was used to test if assumption of sphericity had been violated across trials. If Mauchly’s sphericity assumption was violated, then multivariate tests were reported and Greenhouse-Geisser (ϵ) estimates of sphericity were used to correct degrees of freedom. Posthoc tests with Bonferroni corrections and Pairwise comparisons were applied to significant data. Statistical analysis was performed using IBM SPSS Statistics 25 software licensed to Brown University and graphs were generated with R Studio [27] using ggplot2 [28], FactoMineR [29], and ggcorrplot. For a detailed table of statistic performed in all experiments, please refer to Additional Supplement Item 2.

### Data and Code Availability

All raw and analyzed data supporting the current study is deposited in the Brown University Digital Repository and is freely available online (https://repository.library.brown.edu/studio/). Code for running BEER Run and Data Analysis is freely available via the Kaun Lab github (https://github.com/kaunlab).

## Supporting information

Catalano_et_al_2020_Supplemental Data

## End Matter

### Supplemental Information

Five supplemental figures and four additional supplemental items are included with this manuscript.

### Author Contributions and Notes

Conceptualization, K.R.K, R.A., and U.H., Apparatus design, K.R.K., R.A., U.H.; Software Design, N.M.; Methodology, K.R.K., R.A., N.M. J.L.C.; Data curation and investigation, J.L.C., N.M., S.L.S., and T.B., Data validation, J.L.C. and K.R.K., Data analysis, J.L.C.; Data visualization, J.L.C. and K.R.K.; Writing-Original draft, J.L.C. and K.R.K., Writing-Review & Editing, J.L.C., K.R.K., Supervision, N.M., J.L.C., and K.R.K.; Project administration, K.R.K., Funding acquisition, K.R.K.

The authors declare no conflict of interest.

## Acknowledgments

Thanks to T. Tabachnik (HHMI JRC) for assistance with apparatus design and construction, and P. Polidoro (HHMI JRC) for software design and assistance. Thanks to C. Weaver (HHMI JRC), F. Midgley (HHMI JRC) and R. Svirskas (HHMI JRC) for computational support, S. Moorehead (HHMI JFRC) for organizational support, the Janelia Fly Facility (particularly T. Laverty and K. Hibbard) for help with flies, and D. Rinberg for help with olfactometer design. Thanks to former members of the Heberlein Lab at Janelia Research Campus for their fruitlful suggesstions and discussion regarding the BEER Run apparatus, especially G. Shohat-Ophir (Bar-Ilan University), M. Saver (EMBL), C. Kent (York University) and J. Simon (JRC). Thanks to all of the members of the Kaun Lab for their feedback on the data analysis and writing of this manuscript, especially K. Scaplen, and collaborator J. McGeary. Thanks to Brown University undergraduate F. Muhammed for providing preliminary data which later informed experiments included in this manuscript.

