## Supplementary material for "Behavioral features of motivated response to alcohol in *Drosophila*": Catalano_et_al_2020_Supplemental Data

### Behavioral features of motivated response to alcohol in *Drosophila* Supplemental Information

**Table of Contents** **page**

Title page…....................................................................................................................1
Supplemental Figure 1.............................................................................................… 2-3
Behaviors collected across doses reveal formation of aversive and appetitive phenotypes across trials.
Supplemental Figure 2.............................................................................................… 4-5
Initial behaviors collected expose ethanol dose dependent behaviors across trials.
Supplemental Figure 3.............................................................................................… 6
Behavioral and physiological development of ethanol tolerance across trials
Supplemental Figure 4.............................................................................................… 7-8
Operant task perframe features and behavioral classifiers are dynamic across trials
Supplemental Figure 5.............................................................................................… 9-10
Trial heatmaps and behavioral correlation matrices for perframe features and behavioral classifiers.

Additional Supplemental Item 1.....................................................................................11-12
BEER Run Apparatus, Graphical User Interface (GUI) settings, and list of parts.
Additional Supplemental Item 2 ....................................................................................13-18
Complete list of statistical tests performed on data in all figures and supplemental figures.
Additional Supplemental Item 3 ....................................................................................19
Perframe feature descriptions.
Additional Supplemental Item 4 ....................................................................................20-21
PCA component variance and total-within cluster sum of squares.

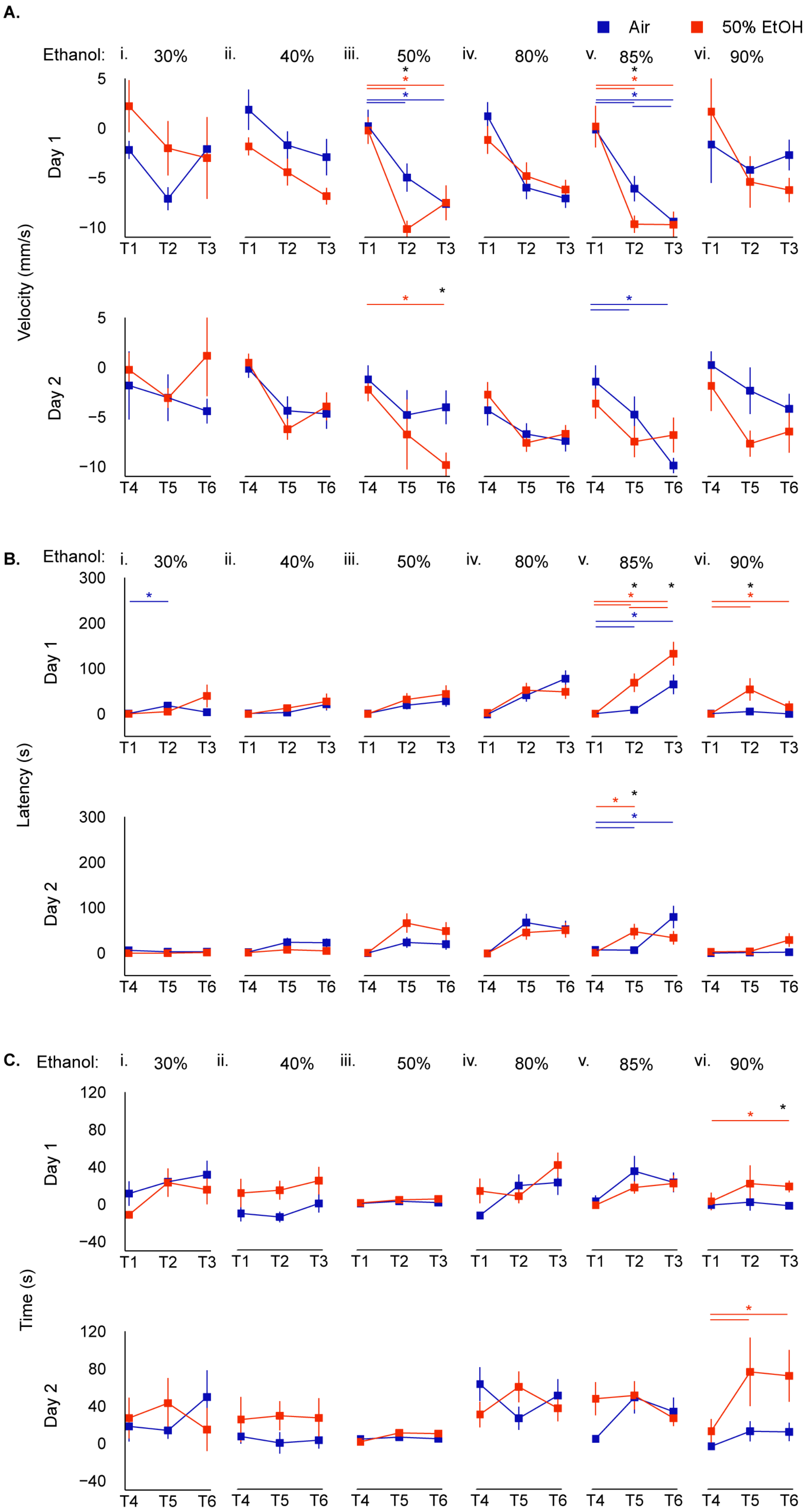
A.
Air
50% EtOH
Ethanol: i. 30% ii. 40% iii. 50% iv. 80% v. 85% vi. 90%
Velocity (mm/s)
Day 1
Day 2
T1 T2 T3
T4 T5 T6
B.
Latency (s)
C.
Time (s)

**Supplementary Figure 1. Behaviors collected across doses reveal formation of aversive and appetitive phenotypes across trials.** Velocity (**A**), latency (**B**), and time (**C**) were recorded for six spaced operant administration trials over two-days at varying concentrations of ethanol *i*. 30% (air n = 5 , ethanol n = 6); *ii*. 40% (air n = 10 , ethanol n = 12); *iii*. 50% (air n = 23 , ethanol n = 19); *iv*. 80 (air n = 23 , ethanol n = 30); *v*. 85% (air n = 22 , ethanol n = 22); *vi*. 90% (air n = 12 , ethanol n = 9). Average behavior during the operant task was collected for each fly in air and ethanol receiving groups and plotted across six trials standardized to trial one behavior. Square-points indicate group mean +/- SE. Repeated ANOVA with planned contrasts were performed for each ethanol concentration (30%, 40%, 50%, 80%, 85%, 90%) and behavior (velocity, latency, time). If Mauchly's test indicated that the assumption of sphericity had been violated, the Greenhouse-Geiser correction was applied to the data. All posthoc analysis was performed with Bonferroni corrections. (**A**) Repeated measures ANOVA indicates significant differences in operant task velocity in the (*Aii*) 40% ethanol group (F(2,40) = 15.40, p = 0.00; trials T1,T3 p=0.003, T4,T5 p=0.000, T4,T6 p=0.004), (*Aiii*) 50% ethanol group (stats previously reported), (*Aiv*) 80% ethanol group (F(1.51,76.99) = 19.42, p = 0.00, Greenhouse Giesser Correction ε=0.76; day 1 v. day 2 p=0.01, T1,T2 p=0.003, T1,T3 p=0.000, T4,T5 p=0.004, T4,T6 p=0.000), and (Av) 85% ethanol group (F(2,84) = 3.20, p = 0.05; air v. ethanol T2 p=0.026, air group T1,T2 p=0.012, T1,T3 p=0.000, T2,T3 p=0.036, T4,T6 p=0.000, T5,T6 p=0.009, ethanol group T1,T2 p=0.000, T1,T3 p=0.000). (**B**) Significant differences in operant task latency. (*Biv*) 80% ethanol group (F(2,102) = 18.36, p = 0.00, p=0.000; trials T1,T2 p=0.003, T1,T3 p=0.000, T4,T5 p=0.001, T4,T6 p=0.000), (*Bv*) 85% ethanol group (F(1.33,55.76) = 3.91, p = 0.04; air v. ethanol day 1 p=0.010, T2, T3, T5 p=0.006, 0.053, 0.024, air T1,T3 p=0.031, T2,T3 p=0.040, T4,T6 p=0.004, T5,T6 p=0.016, ethanol T1,T2 p=0.000, T1,T3 p=0.000, T2,T3 p=0.016, T3,T6 p=0.005, T4,T5 p=0.002), and (*Bvi*) 90% ethanol group (F(1.11,21.17) = 4.73, p = 0.04; air v. ethanol p=0.025, day 1 p=0.026, T2 p=0.027, ethanol T1,T2 p=0.008, T2,T3 p=0.010, T2,T5 p=0.004). (**C**) Significant differences in operant task time. *(Ci)* 40% ethanol group (F(2,40) = 0.002, p = 1.00; air v. ethanol p=0.04), (*Civ*) 80% ethanol group (F(2,102) = 4.93, p = 0.01; air group T1,T4 p=0.001, ethanol group T2,T5 p=0.003), (*Cv*) 85% ethanol group (F(1.68,70.47) = 4.54, p = 0.02; day 1 v. day 2 p=0.008, T1,T2 p=0.027, T1,T3 p=0.015), and (*Cvi*) 90% ethanol group (F(2,38) = 3.77, p = 0.03; air v. ethanol p=0.033, day 2 p=0.046, T3 p=0.003, ethanol group T1,T3 p=0.000,T2,T5 p=0.009, T3,T6 p=0.049, T4,T5 p=0.022, T4,T6 p=0.015). For detailed statistics see Table S2.

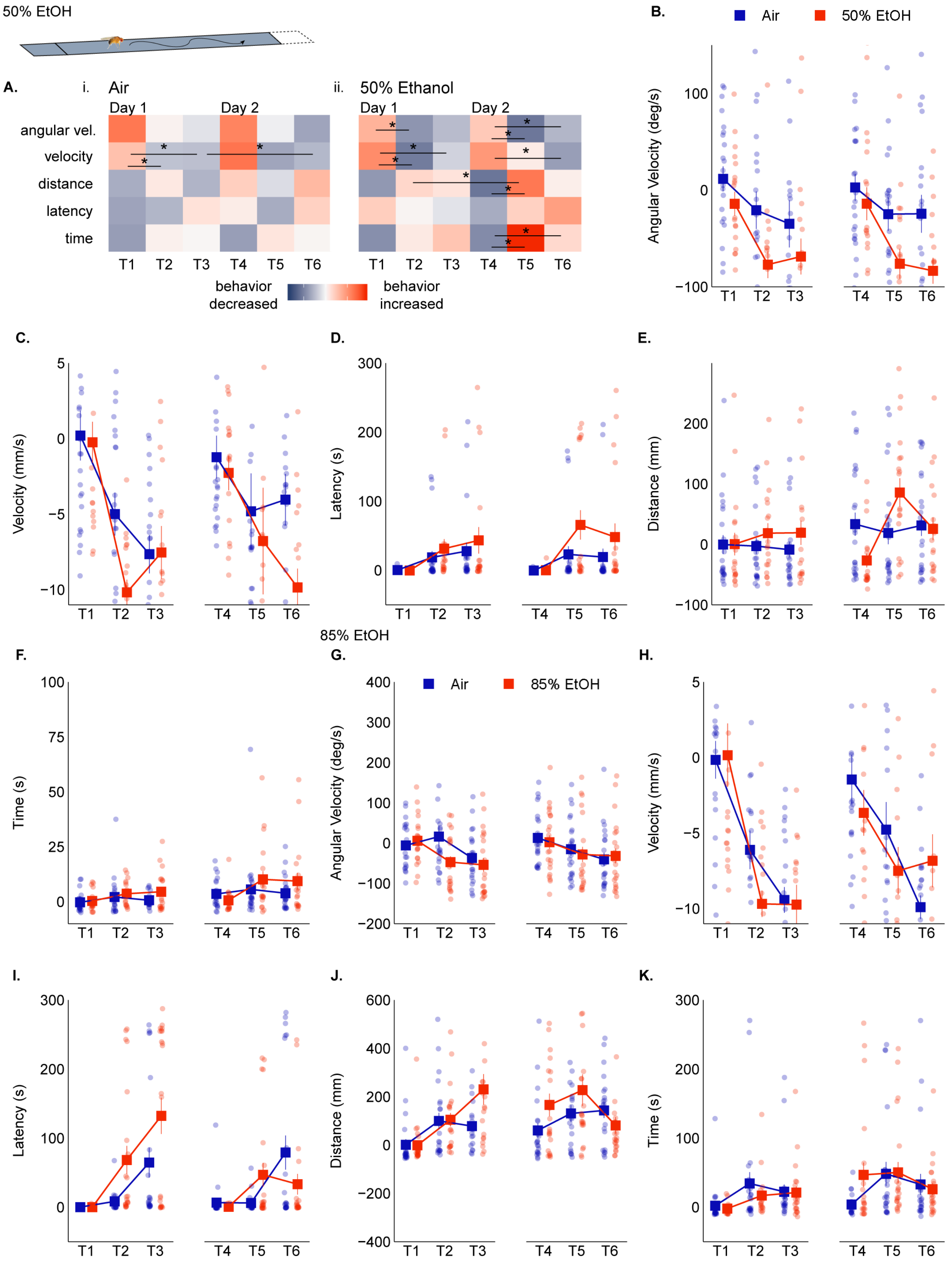
50% EtOH
A.
i. Air
Day 1
Day 2
angular vel.
velocity
distance
latency
time
T1 T2 T3 T4 T5 T6
ii. 50% Ethanol
behavior decreased
behavior increased
B.
Air
50% EtOH
Angular Velocity (deg/s)
C.
Velocity (mm/s)
D.
Latency (s)
E.
Distance (mm)
85% EtOH
F.
Time (s)
G.
85% EtOH
Angular Velocity (deg/s)
H.
Velocity (mm/s)
I.
Latency (s)
J.
Distance (mm)
K.
Time (s)

**Supplementary Figure 2. Initial behaviors collected expose ethanol dose dependent behaviors across trials.** (**A**) Heatmaps of behavioral dynamics for (*i*) air and (*ii*) 50% ethanol groups (air n = 23, ethanol n = 19) for all trials. Rows correspond to a behavior (angular velocity, velocity, distance, latency, time) and columns correspond to trials 1-6. Each grid corresponds to a behavior index calculated as $[\frac{1}{n}\sum_{i=1}^{n}(behavior_{(trial\,y)})_i]$ . Color indicates how much more (red) or less (blue) the behavior occurred in the air or ethanol group. (**B-K**) Behavior counts for angular velocity, velocity, latency, distance, and time were recorded for six spaced operant administration trials over two-days at varying concentrations of ethanol (**B-F** 50%, **G-K** 85%). Average behavior during the operant task was quantified for each fly in the air and ethanol receiving groups and plotted across six trials. Square-points indicate group mean +/- SE. (**B-F**) Repeated measures ANOVA with planned contrasts and posthoc Bonferroni corrections indicate that the 50% ethanol group shows significant differences in operant task behaviors including (A,B), angular velocity (F(2,80) = 12.06, p = 0.00; air v. ethanol p=0.006, day 1, 2 p=0.033, 0.007, T2, T5, T6 p=0.029, 0.036, 0.022, ethanol group T1,T2 p=0.007, T4,T5 p=0.048, T4,T6 p=0.018), (A,C) velocity (MANOVA, Wilks' Lambda day*trials*group: V = 0.86, F(2,39) = 3.16, p =0.05, $\omega^2$ = 0.14; air v. ethanol T2, T6 p=0.005, 0.011, air group T1,T2 p=0.004, T1,T3 p=0.000, T3,T6 p=0.05, ethanol group T1,T2 p=0.000, T1,T3 p=0.002, T4,T6 p=0.004), and (**A,F**) time (MANOVA, Wilks' Lambda trials*group: V = 0.84, F(2,39) = 3.66, p =0.04, $\omega^2$ = 0.16; ethanol group T4,T5 p=0.026, T4,T6 p=0.020). (G-K) Repeated measures ANOVA with planned contrasts and posthoc Bonferroni corrections indicate significant differences in 85% ethanol (air n = 22 , ethanol n = 22) behavioral counts (**G-K**) for (**G**) angular velocity (F(2,84) = 7.95, p = 0.00; trials T1,T3 p=0.004, T4,T6 p=0.026), (**H**) velocity (F(2,84) = 3.20, p = 0.05; air v. ethanol T2 p=0.026, air group T1,T2 p=0.012, T1,T3 p=0.000, T2,T3 p=0.036, T4,T6 p=0.000, T5,T6 p=0.009, ethanol group T1,T2 p=0.000, T1,T3 p=0.000), (**I**) latency (F(1.33,55.76) = 3.91, p = 0.04; air v. ethanol day 1 p=0.010, T2, T3, T5 p=0.006, 0.053, 0.024, air group T1,T3 p=0.031, T2,T3 p=0.040, T4,T6 p=0.004, T5,T6 p=0.016, ethanol group T1,T2 p=0.000, T1,T3 p=0.000, T2,T3 p=0.016, T3,T6 p=0.005, T4,T5 p=0.002), (**K**) time (F(1.68,70.47) = 4.54, p = 0.02; day 1 v. day 2 p=0.008, T1,T2 p=0.027, T1,T3 p=0.015). For detailed statistics see Table S2.

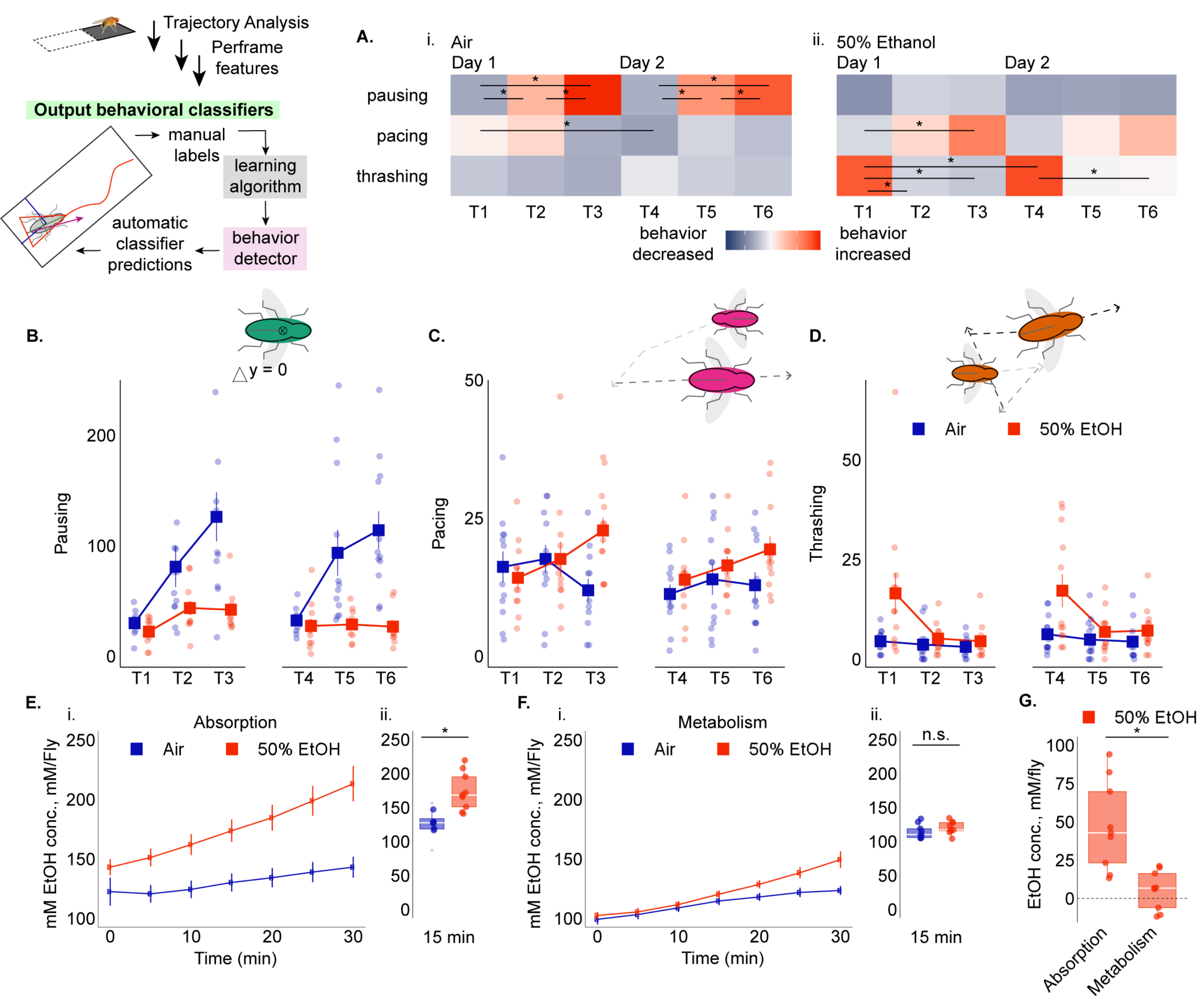


**Supplementary Figure 3. Behavioral and physiological development of ethanol tolerance across trials.** The BEER run operant task paired with detailed trajectory analysis (Ctrax version 0.3.1) and behavioral classification (JAABA) produced behavioral classifier counts for pausing, pacing, thrashing during ethanol administration across six-trials for 24 flies (air n = 12, 50% ethanol n = 12). (**A**) Heatmaps of behavioral dynamics for (*i*) air and (*ii*) 50% ethanol groups for all trials. Rows correspond to a behavior (pausing, pacing, thrashing) and columns correspond to trials 1-6. Each grid corresponds to a behavior index calculated as $[\frac{1}{n}\sum_{i=1}^{n}(behavior_{(trial\,y)})_i]$. Color indicates how much more (red) or less (blue) the behavior occurred in the air or ethanol group. (**B-D**)Behavior counts for pausing, pacing, and thrashing were recorded for six 10min 50% ethanol administrations over two-days. Square-points indicate group mean +/- SE. Repeated measures ANOVA with planned contrasts and posthoc Bonferroni corrections indicate significant differences during administration. The air (**A*i***) and 50% ethanol (**A*ii***) groups show significant differences in (**Ai,B**) pausing (F(2,44) = 14.09, p = 0.00; air v. ethanol p=0.001, day 1, 2 p=0.004, 0.001, T3, T5, T6 p=0.001, 0.006, 0.000, air group T1,T2 p=0.004,T1,T3 p=0.000, T2,T3 p=0.001, T4,T5 p=0.001, T4,T6 p=0.000, T5,T6 p=0.041), (Ai,C) pacing (F(2,44) = 9.56, p = 0.00; air v. ethanol T3 p=0.002, air group T1,T4 p=0.044 , ethanol group T1,T3 p=0.015) and (Ai,D) thrashing (F(1.17,25.74) = 5.77, p = 0.02; air v. ethanol p=0.023, day 1, 2 p=0.033, 0.048, T1,T4 p=0.031, 0.020, ethanol group T1,T2 p=0.005, T1,T3 p=0.016, T4,T5 p=0.002, T4,T6 p=0.002). (**B,D**) Average behaviors (pausing, pacing, thrashing) during administration was quantified for each fly in the air and ethanol receiving groups and plotted across six trials. Square-points indicate group mean +/- SE. (**E**) The 50% ethanol group (n=9) contained higher internal ethanol concentrations than the air control group (n=9) immediately following 1 trial of exposure (absorption) (Levene's Test F(16) = 0.763, p=0.395 (equal variance); Independent Group t-test t(16) = -3.603, p = 0.002). (**F**) The 50% ethanol does not have statistically different internal ethanol concentrations compared to the air control group following 50min metabolism period post-exposure (Levene's Test F(16) = 0.147, p=0.706 395 (equal variance); Independent Group t-test t(16) = -1.214, p = 0.242). (**G**) Flies that received 50% ethanol for 10min had an internal ethanol concentration 42.81 +/- 9.69 mM/fly, and after 50min of rest internal ethanol concentration is a minimal 6.70 +/- 4.22 mM/fly. This significant reduction in internal ethanol concentration (Levene's Test F(16) = 5.215, p=0.036, Independent Group t-test t(16) = 3.976, p = 0.002) suggests near full ethanol metabolism in-between trials. For detailed statistics see Table S2.

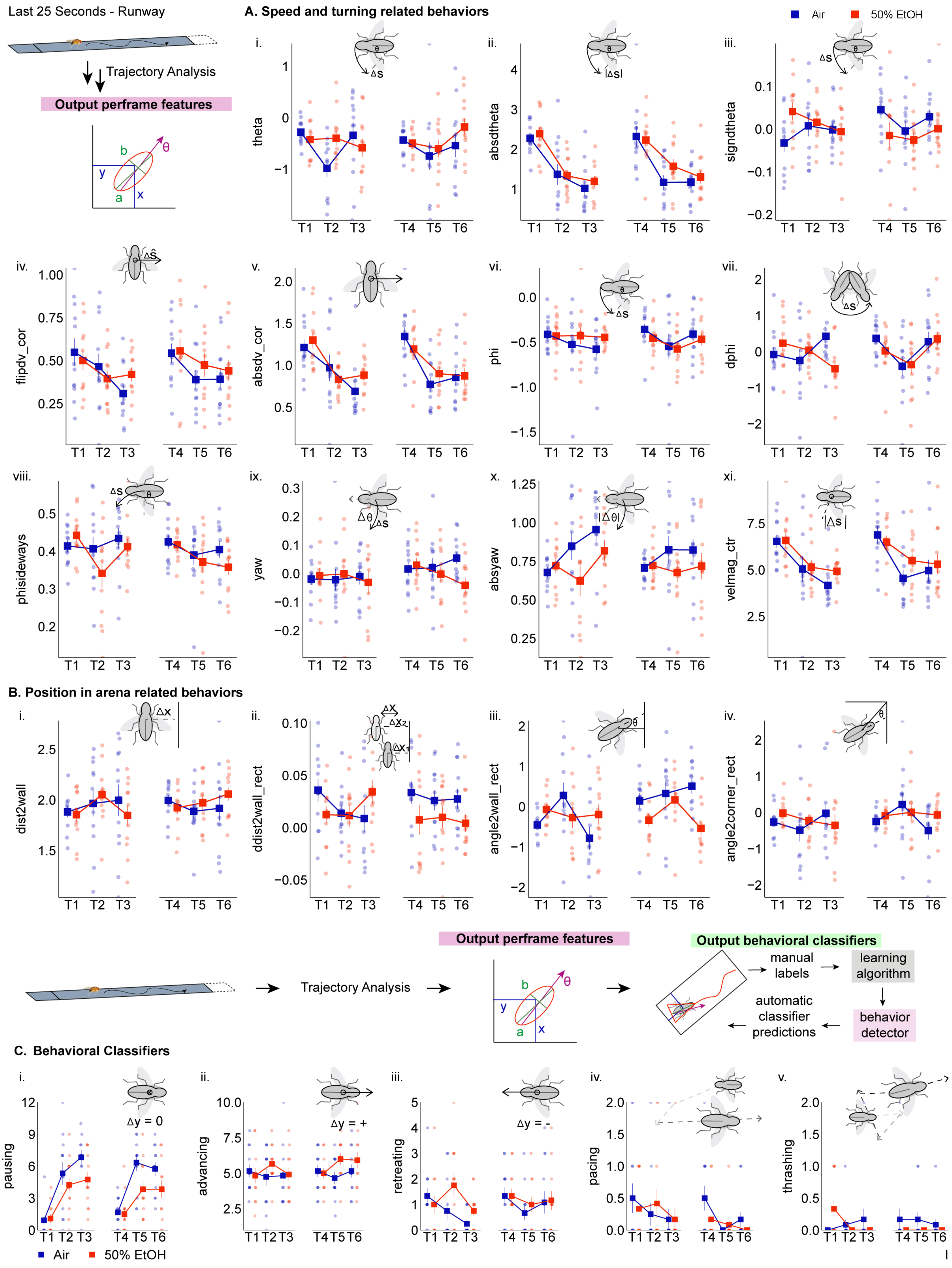

Last 25 Seconds - Runway
Trajectory Analysis
Output perframe features
A. Speed and turning related behaviors
Air
50% EtOH
i.
theta
ii.
absdtheta
iii.
signdtheta
iv.
flipdv_cor
v.
absdv_cor
vi.
phi
vii.
dphi
viii.
phisideways
ix.
yaw
x.
absyaw
xi.
velmag_ctr
T1 T2 T3 T4 T5 T6
B. Position in arena related behaviors
i.
dist2wall
ii.
ddist2wall_rect
iii.
angle2wall_rect
iv.
angle2corner_rect
Output perframe features
Output behavioral classifiers
Trajectory Analysis
manual labels
learning algorithm
automatic classifier predictions
behavior detector
C. Behavioral Classifiers
i.
pausing
Δy = 0
ii.
advancing
Δy = +
iii.
retreating
Δy = -
iv.
pacing
v.
thrashing
Air
50% EtOH

**Supplementary Figure 4. Operant task perframe features and behavioral classifiers are dynamic across trials.** The last 25 seconds of the BEER run operant task was paired with detailed trajectory analysis (Ctrax version 0.3.1) to quantify behavioral pre-frame features during the operant task across six-trials for 24 flies (air n = 12, 50% ethanol n = 12). (**Ai-xi**) Average counts for speed and turning related features (**A**: ***i***. theta, ***ii***. absdtheta, ***iii***. signdtheta, ***iv***. flipdv_cor, ***v***. absdv-cor, ***vi***. phi, *viii*. dphi, *viii*. phisideways, *ix*. yaw, *x*. absyaw, *xi*. velmag). (**B*i-iv***) Average counts for position related features (**B: *i***. dist2wall, ***ii***. ddist2wall, ***iii***. angle2wall, ***iv***. angle2corner) during the operant administration task was quantified for each fly in the air and ethanol receiving groups and plotted across six trials. Square-points indicate group mean +/- SE. Repeated measures ANOVA with planned contrasts and posthoc Bonferroni corrections indicate significant differences in angle2wall (F(2,44) = 4.54, p = 0.02; air v. ethanol day 2 p=0.001, T1, T6 p=0.016, 0.000, air groupT1,T4 p=0.005, T3,T6 p=0.000, T5,T6 p=0.005), phisideways (F(2,44) = 4.96, p = 0.01; T4,T6 p=0.038), absdtheta (F(2,44) = 51.66, p = 0.00; trials T1,T2 p= 0.000, T1,T3 p=0.000, T4,T5 p=0.000, T4,T6 p=0.000), absdv_cor (F(2,44) = 23.02, p = 0.00; trials T1,T2 p= 0.014, T1,T3 p=0.000, T4,T5 p=0.000, T4,T6 p=0.000), signdtheta (F(1,22) = 6.52, p = 0.02; air v. ethanol T1 p=0.01, air group T1,T4 p=0.021), absyaw (MANOVA, Wilks' Lambda trials*group: V = 0.71, F(2,21) = 4.29,  p =0.028, $\omega^2$ = 0.29; T1,T3 p=0.004, air group T1,T3 p=0.002), velmag (F(2,44) = 19.07, p = 0.00; trials T1,T2 p=0.022, T1,T3 p=0.000, T4,T5 p=0.004, T4,T6 p=0.004), flipdv_cor (F(2,44) = 6.15, p = 0.00; T1,T3 p=0.036), dphi (F(2,44) = 4.00, p = 0.03; air v. ethanol T3 p=0.015, air group T4,T5 p=0.044, ethanol group T3,T6 p=0.014), and theta (F(2,44) = 3.36, p = 0.04; air v. ethanol T2 p=0.013, air group T1,T2 p=0.005, T2,T3 p=0.040). (**C**) Average counts for behavioral classifiers obtained through computer vision (Ctrax version 0.3.1) and machine learning (JAABA) pipeline (**C**: ***i***. pausing, ***ii***. advancing, ***iii***. retreating, ***iv***. pacing, ***v***. thrashing) were quantified for the operant task across six-trials for 24 flies (air n = 12, 50% ethanol n = 12). Square-points indicate group mean +/- SE. Repeated measures ANOVA with planned contrasts and posthoc Bonferroni corrections does not indicate significance in advancing (F(2,44) = 3.22, p = 0.05; air v. ethanol T5 p=0.032) and pausing (MANOVA, Wilks' Lambda trials*group: V = 0.97, F(2,21) = 0.25,  p =0.023, $\omega^2$ = 0.30; air v. ethanol p=0.008, day 2 p=0.01, T3, T5 p=0.05, 0.03, air group T1,T2 p=0.003, T1,T3 p=0.000, T4,T5 p=0.000, T4,T6 p=0.000, ethanol group T1,T2 p=0.036, T1,T3 p=0.001, T4,T5 p=0.042, T4,T6 p=0.023). For detailed statistics see Table S2.

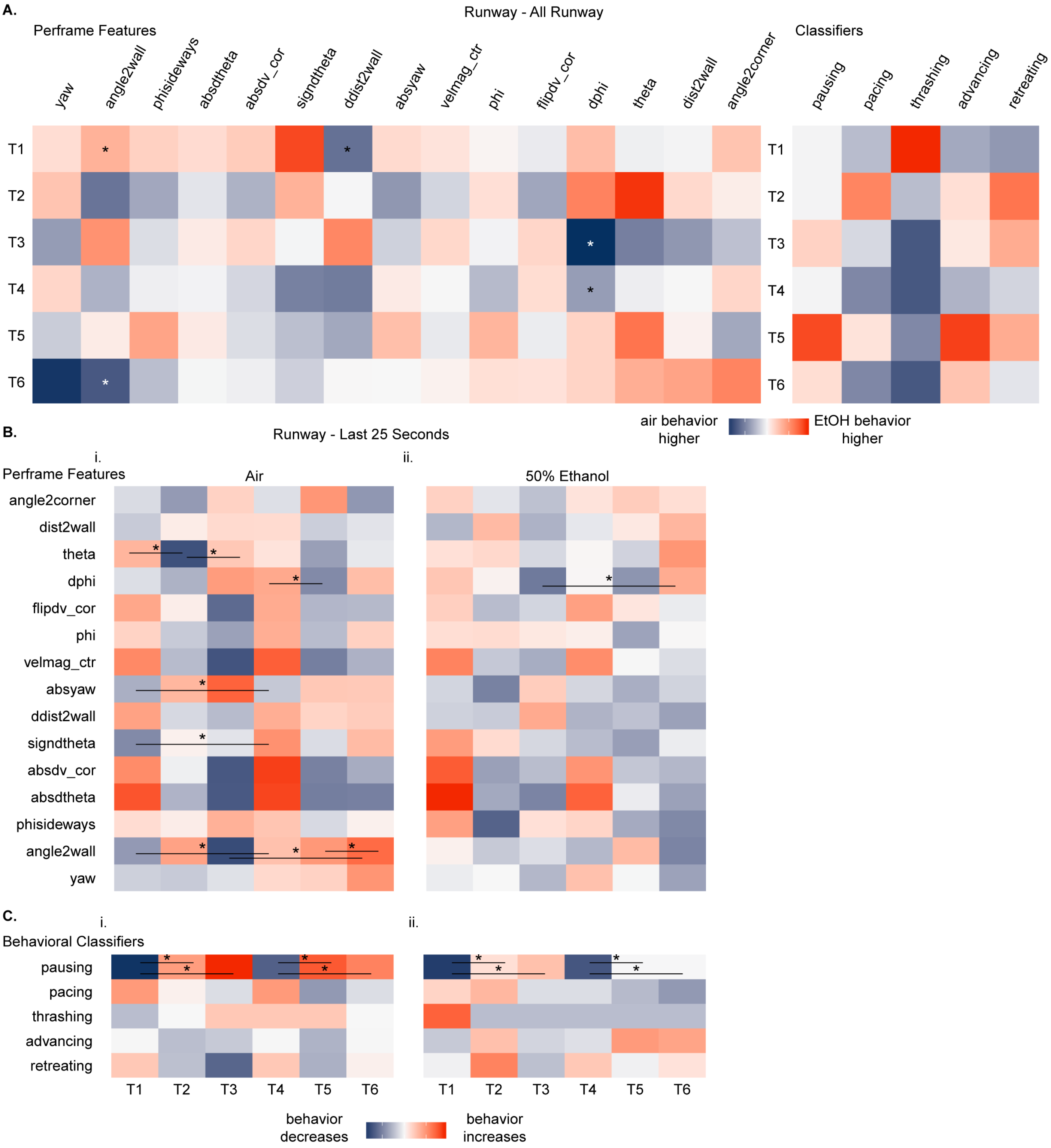
A.
Runway - All Runway
Perframe Features
yaw
angle2wall
phisideways
absdtheta
absdv_cor
signdtheta
ddist2wall
absyaw
velmag_ctr
phi
flipdv_cor
dphi
theta
dist2wall
angle2corner
Classifiers
pausing
pacing
thrashing
advancing
retreating
T1
T2
T3
T4
T5
T6
air behavior higher
EtOH behavior higher
B.
Runway - Last 25 Seconds
i.
ii.
Perframe Features
Air
50% Ethanol
angle2corner
dist2wall
theta
dphi
flipdv_cor
phi
velmag_ctr
absyaw
ddist2wall
signdtheta
absdv_cor
absdtheta
phisideways
angle2wall
yaw
C.
i.
ii.
Behavioral Classifiers
pausing
pacing
thrashing
advancing
retreating
T1
T2
T3
T4
T5
T6
behavior decreases
behavior increases

**Supplementary Figure 5. Trial heatmaps and behavioral correlation matrices for perframe features and behavioral classifiers.** (**A**) Heatmap for the full operant task includes behavioral perframe features: angle2corner, dist2wall, theta, dphi, flipdv_cor, phi, velmag_ctr, absyaw, ddist2wall, signdtheta, absdv_cor, absdtheta, phisideways, angle2wall, yaw, and behavioral classifiers: pausing, pacing, thrashing, advancing, and retreating. Each grid corresponds to a behavior index calculated as previously stated in Fig.1. Color indicates how much more (red) or less (blue) the behavior occurred in the ethanol group compared to the air group. Repeated measures ANOVA with planned contrasts and posthoc Bonferroni corrections indicate significant differences during the full operant task in angle2wall (F(2,44) = 3.60, p = 0.04; air v. ethanol day 2 p=0.014, T1, T6 p=0.015, 0.007, air T1,T2 p=0.046, T1,T4 p=0.012, T2,T3 p=0.008, T3,T6 p=0.009), ddist2wall (F(2,44) = 4.02, p = 0.03; air v. ethanol day 2 p=0.036, T1 p=0.05), and dphi (F(2,44) = 5.78, p = 0.01; air v. ethanol T3, T4 p=0.001, 0.055, air group T2,T3 p=0.012, ethanol group T1,T3 p=0.05, T3,T6 p=0.004). (**B-C**) Heatmaps of behavioral dynamics for (**B*i*, C*i***) air and (**B*ii*, *Cii***) 50% ethanol groups (air n = 12, ethanol n = 12) for all trials. Rows correspond to a behavior (B. perframe features, C. behavioral classifiers) and columns correspond to trials 1-6. Each grid corresponds to a behavior index calculated as $[\frac{1}{n}\sum_{i=1}^{n}(behavior_{(trial\,y)})_i]$ . Color indicates how much more (red) or less (blue) the behavior occurred in the air or ethanol group. Repeated measures ANOVA with planned contrasts and posthoc Bonferroni corrections indicate that the 50% ethanol group indicates significant differences in: (**B**) Perframe features: angle2wall (F(2,44) = 4.54, p = 0.02; air v. ethanol day 2 p=0.001, T1, T6 p=0.016, 0.000, air groupT1,T4 p=0.005, T3,T6 p=0.000, T5,T6 p=0.005), phisideways (F(2,44) = 4.96, p = 0.01; T4,T6 p=0.038), absdtheta (F(2,44) = 51.66, p = 0.00; trials T1,T2 p= 0.000, T1,T3 p=0.000, T4,T5 p=0.000, T4,T6 p=0.000), absdv_cor (F(2,44) = 23.02, p = 0.00; trials T1,T2 p= 0.014, T1,T3 p=0.000, T4,T5 p=0.000, T4,T6 p=0.000), signdtheta (F(1,22) = 6.52, p = 0.02; air v. ethanol T1 p=0.01, air group T1,T4 p=0.021), absyaw (MANOVA, Wilks' Lambda trials*group: V = 0.71, F(2,21) = 4.29, p =0.028, $\omega^2$ = 0.29; T1,T3 p=0.004, air group T1,T3 p=0.002), velmag (F(2,44) = 19.07, p = 0.00; trials T1,T2 p=0.022, T1,T3 p=0.000, T4,T5 p=0.004, T4,T6 p=0.004), flipdv_cor (F(2,44) = 6.15, p = 0.00; T1,T3 p=0.036), dphi (F(2,44) = 4.00, p = 0.03; air v. ethanol T3 p=0.015, air group T4,T5 p=0.044, ethanol group T3,T6 p=0.014), and theta (F(2,44) = 3.36, p = 0.04; air v. ethanol T2 p=0.013, air group T1,T2 p=0.005, T2,T3 p=0.040), and (**C**) Behavioral Classifiers: advancing (F(2,44) = 3.22, p = 0.05; air v. ethanol T5 p=0.032) and pausing (MANOVA, Wilks' Lambda trials*group: V = 0.97, F(2,21) = 0.25, p =0.023, $\omega^2$ = 0.30; air v. ethanol p=0.008, day 2 p=0.01, T3, T5 p=0.05, 0.03, air group T1,T2 p=0.003, T1,T3 p=0.000, T4,T5 p=0.000, T4,T6 p=0.000, ethanol group T1,T2 p=0.036, T1,T3 p=0.001, T4,T5 p=0.042, T4,T6 p=0.023). For detailed statistics see Table S2.

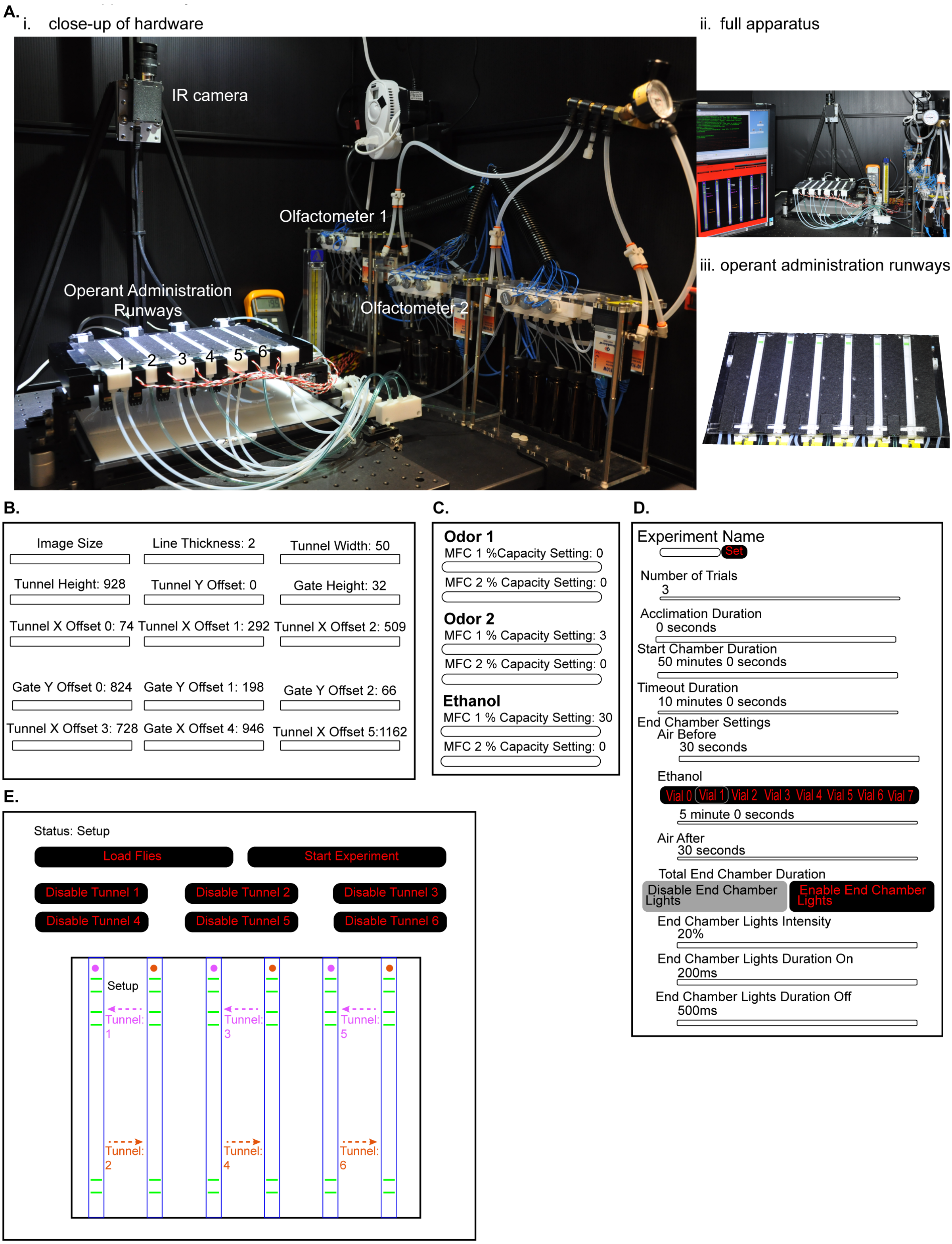


**Additional Supplementary Item 1. BEER Run Apparatus, Graphical User Interface (GUI) settings, and list of parts.** (**A**) Photograph of (***i***) BEER Run operant administration assay consisting of six single-fly runways linked to three olfactometers, and Basler Firewire camera equipped with IR filter, connected to (***ii***) PC running BEER run tracking software, (***iii***) close-up of the 6 operant administration runways. (**B-E**) Representative schematics of the BEER run tracking software GUI, including (**B**) image processing, (**C**) flow rate settings, (**D**) experimental settings, and (**E**) experimental controls. (**F**) Full list of parts used to assemble the BEER Run apparatus.

Additional Supplementary Item 1. (**F**) List of parts for apparatus construction

| Component | Part Number | Vendor |
|---|---|---|
| Runway Arena | 12cm long teflon tube | White Glove Machining Inc. |
| Gates | Custom made gates | 3D printed inhouse |
| Metal Gear Micro Servo | HS_5056MG | Hitec |
| Arena holder | 24cm x 30cm aluminum frame | Inhouse machine shop |
| EL-FLOW (mass flowmeter) | IQF-200C-AAD-00-V-S - with 100 sccm | Bronkhorst USA Inc |
| EL-FLOW (mass flowmeter) | IQF-200C-AAD-00-V-S - with 1000 sccm | Bronkhorst USA Inc |
| Inline 5X2-way Normally close | 161T122 | NResearch Inc. |
| PTFE tubing 1/6" ID | EW-06605-27 | Cole-Parmer |
| 3.5-inch x 2.5-inch, 18 tapped 1/4-20 holes | MX3TA | Siskiyou Corp |
| building block, 1.0 in | BB-1.0 | Siskiyou Corp |
| building block, 5.0 in | BB-5.0 | Siskiyou Corp |
| Water cooled breadboard (12in X 12in) | MBC12 | Thorlabs |
| Breadboard 18x24x1/2 | MB1824 | Thorlabs |
| Adapter Plate for DT12 Stage | DT12B | Thorlabs |
| 1/2" (12mm) Dovetail Translation Stage | DT12 | Thorlabs |
| Rail Carrier, Extended 1 in x 2 in | RC2 | Thorlabs |
| Rail Carrier, Perpendicular Dovetail | RC3 | Thorlabs |
| IR back light illumination | IR 32cm x 32cm | Advanced Illumination |
| PCI Express Card – Dual Bus | FWB-PCIE-02 | Point Grey Red Inc |
| 6-pin to 9-pin, Locking IEEE-1394a to 1394b Cable | ACC-01-2007 | Point Grey Red Inc |
| Basler Firewire Mono Cam | A622FM | Edmund Optics Inc |
| 25MM FL 1” Format Lens | 63246 | Edmund Optics Inc |
| Filter IR MTD 49 X 0.75MM | 54752 | Edmund Optics Inc |
| A600/A622 Mounting Adapter | 58794 | Edmund Optics Inc |

#### Additional Supplementary Item 2. Table of Statistics.

| | Figure | Data Structure, n-value | Statistical Test | Post Hoc |
|---|---|---|---|---|
| | | | 1.) Mauchly's Test of Sphericity. If sphericity is violated (sig. < 0.05) then Greenhouse-Geisser correction (ε) applied to 2.) Repeated Measures Anova | Bonferroni correction w/ pairwise comparisons |
| 1 | 1E.i. 30% ethanol (air n=5 , ethanol n=6) | **Runway - Angular Velocity** Group mean +/- standard error with individuals. All values standardized to day 1, trial 1 data. Data plotted for 6 trials | day*trials*group: $\chi2(2) = 3.85$, $p = 0.15$ $F(2,18) = 4.08$, $p = 0.03$ | #1 (air v. ethanol) T3 p=0.011, T4 p=0.047 #2 (air) T2,T3 p=0.039, T3,T6 p=0.049 #3 (ethanol) T1,T4 p=0.031 |
| 2 | 1E.ii. 40% ethanol (angular velocity) (air n=10, ethanol n=12) | | day*trials: $\chi2(2) = 0.38$, $p = 0.83$ $F(2,40) = 4.73$, $p = 0.01$ | T1,T3 p=0.031, T4,T5 p=0.001, T4,T6 p=0.003 |
| 3 | 1Eiii. 1F. 1Gi. Fig.S2A. Fig.S2B. 50% ethanol (air n=23, ethanol n=19) | | trials: $\chi2(2) = 2.18$, $p = 0.34$ $F(2,80) = 12.06$, $p = 0.00$ test of between-subjects group effect: $F(1,40) = 8.61$, $p = 0.01$ | #1 (air v. ethanol) p=0.006, day 1 p=0.033, day 2 p=0.007, T2 p=0.029, T5 p=0.036, T6 p=0.022 #3 (ethanol) T1,T2 p=0.007, T4,T5 p=0.048, T4,T6 p=0.018 |
| 4 | 1E.iv. 80% ethanol (air n=23, ethanol n=30) | | day*trials*group: $\chi2(2) = 6.98$, $p = 0.03$, ε=0.89 $F(1.77,90.24) = 0.18$, $p = 0.81$ | NA |
| 5 | 1E.v. 85% ethanol (air n=22, ethanol n=22) | | trials: $\chi2(2) = 0.24$, $p = 0.89$ $F(2,84) = 7.95$, $p = 0.00$ | T1,T3 p=0.004, T4,T6 p=0.026 |
| 6 | 1E.vi. 90% ethanol (air n=12, ethanol n= 9) | | day*trials*group: $\chi2(2) = 0.41$, $p = 0.82$ $F(2,38) = 5.73$, $p = 0.01$ | #1 (air v. ethanol) day 1 p=0.048, T1 p=0.012, T2 p=0.005, T6 p=0.015 #2 (air) T1,T2 p=0.004 #3 (ethanol) T2,T3 p=0.028 |
| 7 | 1F. 1Gii. Fig.S2A. Fig.S2C. (velocity) | **Runway** (air n = 23 , 50% ethanol n = 19) Group mean +/- standard error with individuals. All values standardized to day 1, trial 1 data. Data plotted for 6 trials. | trials: $\chi2(2) = 5.66$, $p = 0.06$ $F(2,80) = 20.52$, $p = 0.00$ MANOVA, Wilks' Lambda day*trials*group: $V = 0.86$, $F(2,39) = 3.16$, $p = 0.05$, $\omega^2 = 0.14$ | T1,T2 p=0.000, T1,T3 p=0.000, T4,T6 p=0.003 #1 (air v. ethanol) T2 p=0.005, T6 p=0.011 #2 (air) T1,T2 p=0.004, T1,T3 p=0.000, T3,T6 p=0.05 #3 (ethanol) T1,T2 p=0.000, T1,T3 p=0.002, T4,T6 p=0.004 |
| 8 | 1F. Fig.S2A. Fig.S2A. Fig.S2D. (latency) | | trials: $\chi2(2) = 0.39$, $p = 0.82$ $F(2,80) = 9.28$, $p = 0.00$ | T1,T2 p=0.010, T1,T3 p=0.007, T4,T5 p=0.001, T4,T6 p=0.011 |
| 9 | 1F. Fig.S2A. Fig.S2E. (distance) | | trials*group: $\chi2(2) = 0.62$, $p = 0.73$ $F(2,80) = 4.73$, $p = 0.01$ | day 1, day 2 p=0.022 #1 (air v, ethanol) T4 p=0.014, T5 p=0.029 #3 (ethanol) T2,T5 p=0.018, T4,T5 p=0.000 |
| 10 | 1F. Fig.S2A. Fig.S2F. (time) | | trials: $\chi2(2) = 5.37$, $p = 0.07$ $F(2,80) = 5.68$, $p = 0.01$ MANOVA, Wilks' Lambda trials*group: $V = 0.84$, $F(2,39) = 3.66$, $p = 0.04$, $\omega^2 = 0.16$ | day 1 v. day 2 p=0.005, T4,T5 p=0.05 #3 (ethanol) T4,T5 p=0.026, T4,T6 p=0.020 |
| 11 | 2A. velocity, trial 1 | **Administration - Velocity** (air n = 23 , 50% ethanol n = 19) Group mean binned to 15 seconds +/- standard error plotted against time. Data plotted for 6 trials. | trials*group: $\chi2(2) = 11.95$, $p = 0.00$, ε=0.79 $F(1.58,63.30) = 28.18$, $p = 0.00$ | #1 (air v. ethanol) T1 p=0.000, T4 p=0.003, T6 p=0.027 #2 (air) T1,T2 p=0.000, T1,T3 p=0.000, T2,T3 p=0.000, T4,T5 p=0.000, T4,T6 p=0.000, T5,T6 p=0.000 #3 (ethanol) T1,T2 p=0.000, T1,T3 p=0.000, T4,T5 p=0.019, T4,T6 p=0.005 |
| 12 | 2B. velocity, trial 2 | | | |
| 13 | 2C. velocity, trial 3 | | | |
| 14 | 2D. velocity, trial 4 | | | |
| 15 | 2E. velocity, trial 5 | | | |
| 16 | 2F. velocity, trial 6 | | | |
| 17 | 2H. 2Ii. Fig.S3A. Fig.S3B. (pausing) | **Administration - Classifiers** (air n = 12, 50% ethanol n = 12) Group mean +/- standard error with individuals. Data plotted for 6 trials. | trials*group: $\chi2(2) = 1.09$, $p = 0.58$ $F(2,44) = 14.09$, $p = 0.00$ test of between-subjects group effect: $F(1,22) = 15.82$, $p = 0.00$ | #1 (air v. ethanol) p=0.001, day 1 p=0.004, day 2 p=0.001, T3 p=0.001, T5 p=0.006, T6 p=0.000 #2 (air) T1,T2 p=0.004,T1,T3 p=0.000, T2,T3 p=0.001, T4,T5 p=0.001, T4,T6 p=0.000, T5,T6 p=0.041 |
| 18 | 2H. 2Iii. Fig.S3A. Fig.S3C. (pacing) | | trials*group: $\chi2(2) = 1.21$, $p = 0.55$ $F(2,44) = 9.56$, $p = 0.00$ | #1 (air v. ethanol) T3 p=0.002 #2 (air) T1,T4 p=0.044 #3 (ethanol) T1,T3 p=0.015 |

Table 2b. Statistics Table

| | Figure | Data Structure, n-value | Statistical Test | Post Hoc |
|---|---|---|---|---|
| | | | 1.) Mauchly's Test of Sphericity. If sphericity is violated (sig. < 0.05) then Greenhouse-Geisser correction (ε) applied to 2.) Repeated Measures Anova | Bonferroni correction w/ pairwise comparisons |
| 19 | 2H. 2Iiii. Fig.S3A. Fig.S3D. (thrashing) | **Administration - Classifiers** (air n = 12, 50% ethanol n = 12) Group mean +/- standard error with individuals. Data plotted for 6 trials. | trials*group: χ2(2) = 14.01, p = 0.00, ε=0.67 F(1.17,25.74) = 5.77, p = 0.02 test of between-subjects group effect: F(1,22) = 5.93, p = 0.02 | #1 (air v. ethanol) p=0.023, day 1 p=0.033, day 2 p=0.048, T1 p=0.031, T4 p=0.020 #3 (ethanol) T1,T2 p=0.005, T1,T3 p=0.016, T4,T5 p=0.002, T4,T6 p=0.002 |
| 20 | Fig 3B. yaw | | χ2(2) = 4.28, p = 0.12 F(2,44) = 0.31, p = 0.74 | NA |
| 21 | angle2wall | | trials*day*group: χ2(2) = 1.48, p = 0.48 F(2,44) = 4.54, p = 0.02 | #1 (air v. ethanol) day 2 p=0.001, T1 p=0.016, T6 p=0.000 #2 (air) T1,T4 p=0.005, T3,T6 p=0.000, T5,T6 p=0.005 |
| 22 | phisideways | | trials: χ2(2) = 4.99, p = 0.08 F(2,44) = 4.96, p = 0.01 | T4,T6 p=0.038 |
| 23 | absdtheta | | trials: χ2(2) = 2.30, p = 0.32 F(2,44) = 51.66, p = 0.00 | T1,T2 p= 0.000, T1,T3 p=0.000, T4,T5 p=0.000, T4,T6 p=0.000 |
| 24 | absdv_cor | | trials: χ2(2) = 2.70, p = 0.26 F(2,44) = 23.02, p = 0.00 | T1,T2 p= 0.014, T1,T3 p=0.000, T4,T5 p=0.000, T4,T6 p=0.000 |
| 25 | signdtheta | | days*group: χ2(2) = 0.63, p = 0.73 F(1,22) = 6.52, p = 0.02 | #1 (air v. ethanol) T1 p=0.019 #2 (air) T1,T4 p=0.021 |
| 26 | ddist2wall | | trials*day*group: χ2(2) = 0.67, p = 0.72 F(2,44) = 1.63, p = 0.21 | NA |
| 27 | absyaw | **Last 25 seconds - Runway perframe features** (air n=12, 50% ethanol n=12) z-scores for each behavior plotted in a heatmap | trials: χ2(2) = 5.10, p = 0.08 F(2,44) = 3.47, p = 0.04 MANOVA, Wilks' Lambda trials*group: V = 0.71, F(2,21) = 4.29, p =0.028, $\omega^2$ = 0.29 | T1,T3 p=0.004 #2 (air) T1,T3 p=0.002 |
| 28 | velmag_ctr | | trials: χ2(2) = 1.38, p = 0.50 F(2,44) = 19.07, p = 0.00 | T1,T2 p=0.022, T1,T3 p=0.000, T4,T5 p=0.004, T4,T6 p=0.004 |
| 29 | phi | | day*trials*group: χ2(2) = 0.54, p = 0.76 F(2,44) = 0.12, p = 0.88 | NA |
| 30 | flipdv_cor | | trials: χ2(2) = 0.56, p = 0.76 F(2,44) = 6.15, p = 0.00 | T1,T3 p=0.036 |
| 31 | dphi | | trials*day*group: χ2(2) = 0.87, p = 0.65 F(2,44) = 4.00, p = 0.03 | #1 (air v. ethanol) T3 p=0.015 #2 (air) T4,T5 p=0.044 #3 (ethanol) T3,T6 p=0.014 |
| 32 | theta | | trials*group χ2(2) = 4.71, p = 0.10 F(2,44) = 3.36, p = 0.04 | #1 (air v. ethanol) T2 p=0.013 #2 (air) T1,T2 p=0.005, T2,T3 p=0.040 |
| 33 | dist2wall | | trials*day*group: χ2(2) = 3.01, p = 0.22 F(2,44) = 0.94, p = 0.40 | NA |
| 34 | angle2corner | | days*trials: χ2(2) = 6.68, p = 0.04, ε=0.79 F(1.57,34.58) = 2.59, p = 0.11 | NA |
| 35 | Fig 3B. thrashing | | χ2(2) = 11.31, p = 0.00, ε=0.71 F(1.41,31.06) = 3.55, p = 0.056 | NA |
| 36 | pacing | | trials: χ2(2) = 2.62, p = 0.27 F(2,44)= 3.51, p = 0.039 | NA |
| 37 | retreating | | day*trials*group: χ2(2) = 4.45, p = 0.11 F(2,44) = 0.65, p = 0.53 | NA |
| 38 | advancing | **Last 25 seconds - Runway classifier** (air n=12, 50% ethanol n=12) z-scores for each behavior plotted in a heatmap | trials*group: χ2(2) = 0.64, p = 0.73 F(2,44) = 3.22, p = 0.05 | #1 (air v. ethanol) T5 p=0.032 |
| 39 | pausing | | trials: χ2(2) = 7.39, p = 0.03, ε=0.77 F(1.54,33.93) = 36.51, p = 0.00 MANOVA, Wilks' Lambda trials*group: V = 0.97, F(2,21) = 0.25, p =0.023, $\omega^2$ = 0.30 | #1 (air v. ethanol) p=0.008, day 2 p=0.01, T3 p=0.05, T5 p=0.03 #2 (air) T1,T2 p=0.003, T1,T3 p=0.000, T4,T5 p=0.000, T4,T6 p=0.000 #3 (ethanol) T1,T2 p=0.036, T1,T3 p=0.001, T4,T5 p=0.042, T4,T6 p=0.023 |

Table 2c. Statistics Table

| | Figure | Data Structure, n-value | Statistical Test<br>1.) Mauchly's Test of Sphericity. If sphericity is violated (sig. < 0.05) then Greenhouse-Geisser correction (ε) applied to 2.) Repeated Measures Anova | Post Hoc<br>Bonferroni correction w/ pairwise comparisons |
|---|---|---|---|---|
| 40 | Fig 3C.<br>yaw | **Just Latency - Runway perframe features**<br>(air n=12, 50% ethanol n=12) z-scores for each behavior plotted in a heatmap | day*trials*group: χ2(2) = 0.54, p = 0.76<br>F(2,44) = 0.34, p = 0.71 | NA |
| 41 | angle2wall_rect | | day*trials*group: χ2(2) = 2.57, p = 0.28<br>F(2,44) = 6.18, p = 0.00 | #1 (air v. ethanol) T2 p=0.030, T3 p=0.055, T6 p=0.010 #2 (air) T1,T2 p=0.017, T2,T3 p=0.003 |
| 42 | phisideways | | day*trials*group: χ2(2) = 7.17, p = 0.03, ε=0.78<br>F(1.55,34.13) = 0.85, p = 0.41 | NA |
| 43 | absdtheta | | days*group: χ2(2) = 7.04, p = 0.03, ε=0.78<br>F(1,22) = 4.36, p = 0.05<br>trials: F(1.70,37.44) = 74.72, p = 0.00 | T1,T2 p=0.000, T1,T3 p=0.000, T4,T5 p=0.000, T4,6 p=0.000 #1 (air v. ethanol) T6 p=0.032 |
| 44 | absdv_cor | | days*group: χ2(2) = 10.26, p = 0.01, ε=0.72<br>F(1,22) = 9.67, p = 0.01<br>trials: F(1.79,39.46) = 39.99, p = 0.00 | T1,T2 p=0.000, T1,T3 p=0.000, T4,T5 p=0.006, T4,6 p=0.000 #1 (air v. ethanol) day 1 p=0.016, T1 p=0.049, T6 p=0.036 |
| 45 | signdtheta | | day*trials*group: χ2(2) = 6.29, p = 0.04, ε=0.79<br>F(1.59,34.96) = 4.11, p = 0.03 | #1 (air v. ethanol) day 2 p=0.024, T1 p=0.019, T3 p=0.038 #2 (air) T1,T4 p=0.035 |
| 46 | ddist2wall | | day*trials*group: χ2(2) = 8.60, p = 0.01, ε=0.75<br>F(1.50,32.94) = 0.78, p = 0.43 | NA |
| 47 | absyaw | | trials: χ2(2) = 2.08, p = 0.35<br>F(2,44) = 12.15, p = 0.00 | T1,T3 p=0.002, T4,T6 p=0.002 |
| 48 | velmag_ctr | | χ2(2) = 4.09, p = 0.13<br>day*group: F(1,22) = 5.38, p = 0.03<br>trials: F(2,44) = 76.88, p = 0.00 | T1,T2 p=0.000, T1,T3 p=0.000, T4,T5 p=0.000, T4,T6 p=0.000 #1 (air v. ethanol) T5 p=0.035 |
| 49 | phi | | day*trials*group: χ2(2) = 2.38, p = 0.30<br>F(2,44) = 0.76, p = 0.47 | NA |
| 50 | flipdv_cor | | trials: χ2(2) = 4.04, p = 0.13<br>F(2,44) = 5.10, p = 0.01 | T1 v. T3 p=0.015 |
| 51 | dphi | | day*trials*group: χ2(2) = 0.95, p = 0.62<br>F(2,44) = 3.44, p = 0.04 | NA |
| 52 | theta | | day*trials*group: χ2(2) = 0.49, p = 0.79<br>F(2,44) = 2.70, p = 0.08 | NA |
| 53 | dist2wall | | day*trials*group: χ2(2) = 3.43, p = 0.18<br>F(2,44) = 0.87, p = 0.43 | NA |
| 54 | angle2corner | | day*trials*group: χ2(2) = 2.77, p = 0.25<br>F(2,44) = 1.99, p = 0.15 | NA |
| 55 | Fig 3C.<br>thrashing | **Just Latency - Runway classifier**<br>(air n=12, 50% ethanol n=12) z-scores for each behavior plotted in a heatmap | day*trials*group: χ2(2) = 10.49, p = 0.01, ε=0.72<br>F(1.44,31.58) = 3.49, p = 0.06 | NA |
| 56 | pacing | | day: χ2(2) = 11.18, p = 0.01, ε=0.71<br>F(1,22) = 5.06, p = 0.04 | Day 1 v. day 2 p=0.035 |
| 57 | retreating | | day*trials*group: χ2(2) = 4.25, p = 0.12<br>F(2,44) = 0.91, p = 0.41 | NA |
| 58 | advancing | | trials: χ2(2) = 1.35, p = 0.51<br>F(2,44) = 1.51, p = 0.02 | T1,T3 p=0.035 |
| 59 | pausing | | days*trials: χ2(2) = 5.00, p = 0.08<br>F(2,44) = 3.22, p = 0.05 | T1,T2 p=0.05, T4,T5 p=0.007 |

Table 2d. Statistics Table

| | Figure | Data Structure, n-value | Statistical Test | Post Hoc |
|---|---|---|---|---|
| | | | Paired T-test | |
| 60 | 4H. Behaviors that increase with training | Runway (air n = 12, 50% ethanol n = 12) Behavior value = Ethanol – Air group mean | (angular_velocity) T2,T3 t(11) = -1.99, p = 0.07; T5,T6 t(11) = -0.21, p = 0.84<br>(pausing) T2,T3 t(11) = 1.10, p = 0.30; T5,T6 t(11) = -0.44, p = 0.67<br>(theta) T2,T3 t(11) = 3.13, p = 0.01; T5,T6 t(11) = -1.22, p = 0.25<br>(absyaw) T2,T3 t(11) = -0.77, p = 0.46; T5,T6 t(11) = -0.50, p = 0.63 | |
| 61 | 4I. Behaviors that decrease with training | | (velocity) T2,T3 t(11) = -1.71, p = 0.12; T5,T6 t(11) = 1.12, p = 0.29<br>(absdtheta) T2,T3 t(11) = -1.04, p = 0.32; T5,T6 t(11) = -0.50, p = 0.63<br>(angle2wall) T2,T3 t(11) = -4.12, p = 0.00; T5,T6 t(11) = 3.46, p = 0.01<br>(latency) T2,T3 t(11) = -0.59, p = 0.57; T5,T6 t(11) = 1.32, p = 0.22<br>(advancing) T2,T3 t(11) = 1.41, p = 0.19; T5,T6 t(11) = 1.05, p = 0.32 | |
| | | | 1.) Mauchly's Test of Sphericity. If sphericity is violated (sig. < 0.05) then Greenhouse-Geisser correction (ε) applied to 2.) Repeated Measures Anova | Bonferroni correction w/ pairwise comparisons |
| 62 | Fig.S1Ai.<br>30% ethanol<br>(velocity)<br>(air n=5 , EtOH n= 6) | Runway Group mean +/- standard error with individuals. All values standardized to day 1, trial 1 data. Data plotted for 6 trials | day*trials*group: χ2(2) = 1.11, p = 0.57<br>F(2,18) = 4.30, p = 0.03 | NA |
| 63 | Fig.S1Aii.<br>40% ethanol<br>(velocity)<br>(air n=10 , EtOH n=12) | | trials: χ2(2) = 3.08, p = 0.21<br>F(2,40) = 15.40, p = 0.00 | T1,T3 p=0.003, T4,T5 p=0.000, T4,T6 p=0.004 |
| 64 | Fig.S1Aiv.<br>80% ethanol<br>(velocity)<br>(air n=23 , EtOH n=30) | | trials: χ2(2) = 19.65, p = 0.00, ε=0.76<br>F(1.51,76.99) = 19.42, p = 0.00 | day 1 v. day 2 p=0.01, T1,T2 p=0.003, T1,T3 p=0.000, T4,T5 p=0.004, T4,T6 p=0.000 |
| 65 | Fig.S1Avi.<br>90% ethanol<br>(velocity)<br>(air n=12 , EtOH n=9) | | trials: χ2(2) = 10.39, p = 0.01, ε=0.70<br>F(1.39,26.42) = 6.41, p = 0.01 | NA |
| 66 | Fig.S1Bi.<br>30% ethanol<br>(latency)<br>(air n=5 , EtOH n= 6) | Runway Group mean +/- standard error with individuals. All values standardized to day 1, trial 1 data. Data plotted for 6 trials | day*trials*group: χ2(2) = 9.82, p = 0.01, ε=0.59<br>F(1.17,10.55) = 3.37, p = 0.09 | #1 (air v. ethanol) day 2 p=0.040 #2 (air) T1,T2 p=0.021, T1,T4 p=0.05, T2,T5 p=0.034 #3 (ethanol) T3,T6 p=0.041 |
| 67 | Fig.S1Bii.<br>40% ethanol<br>(latency)<br>(air n=10 , EtOH n=12) | | trials: χ2(df) = 11.53, p = 0.00, ε=0.69<br>F(1.38,27.49) = 4.13, p = 0.04 | T4,T6 p=0.018 |
| 68 | Fig.S1Biv.<br>80% ethanol<br>(latency)<br>(air n=23 , EtOH n=30) | | trials: χ2(2) = 4.27, p = 0.12<br>F(2,102) = 18.36, p = 0.00 | T1,T2 p=0.003, T1,T3 p=0.000, T4,T5 p=0.001, T4,T6 p=0.000 |
| 69 | Fig.S1Bvi.<br>90% ethanol<br>(latency)<br>(air n=12 , EtOH n=9) | | day*trials*group: χ2(2) = 28.52, p = 0.00, ε=0.56<br>F(1.11,21.17) = 4.73, p = 0.04 | #1 (air v. ethanol) p=0.025, day 1 p=0.026, T2 p=0.027 #3 (ethanol) T1,T2 p=0.008, T2,T3 p=0.010, T2,T5 p=0.004 |
| 70 | Fig.S1Ci.<br>30% ethanol<br>(time)<br>(air n=5 , EtOH n= 6) | Runway Group mean +/- standard error with individuals. All values standardized to day 1, trial 1 data. Data plotted for 6 trials | day*trials*group: χ2(2) = 10.45, p = 0.01, ε=0.58<br>F(1.16,10.41) = 0.79, p = 0.41 | NA |
| 71 | Fig.S1Cii.<br>40% ethanol<br>(time)<br>(air n=10 , EtOH n=12) | | day*trials*group: χ2(2) = 0.59, p = 0.75<br>F(2,40) = 0.002, p = 1.00<br>test of between-subject group effects: F(1,20) = 4.79, p = 0.04 | #1 (air v. ethanol) p=0.04 |

Table 2e. Statistics Table

| | Figure | Data Structure, n-value | Statistical Test | Post Hoc |
|---|---|---|---|---|
| | | | 1.) Mauchly's Test of Sphericity. If sphericity is violated (sig. < 0.05) then Greenhouse-Geisser correction (ε) applied to 2.) Repeated Measures Anova | Bonferroni correction w/ pairwise comparisons |
| 72 | Fig.S1Civ. 80% ethanol (time) (air n=23 , EtOH n=30) | **Runway** Group mean +/- standard error with individuals. All values standardized to day 1, trial 1 data. Data plotted for 6 trials | day*trials*group χ2(2) = 1.08, p = 0.58 F(2,102) = 4.93, p = 0.01 | day 1 v. day 2 p=0.000 #2 (air) T1,T4 p=0.001 #3 (ethanol) T2,T5 p=0.003 |
| 73 | Fig.S1Cvi. 90% ethanol (time) (air n=12 , EtOH n=9) | | trials*group: χ2(2) =1.50, p = 0.47 F(2,38) = 3.77, p = 0.03 | day 1 v. day 2 p=0.019 #1 (air v. ethanol) p=0.033, day 2 p=0.046, T3 p=0.003 #3 (ethanol) T1,T3 p=0.000,T2,T5 p=0.009, T3,T6 p=0.049, T4,T5 p=0.022, T4,T6 p=0.015 |
| 74 | Fig.S2Av. Fig.S2G (velocity) | **85% Ethanol - Runway** (air n=22, 85% ethanol n=22) Group mean +/- standard error with individuals. All values standardized to day 1, trial 1 data. Data plotted for 6 trials | χ2(2) = 5.31, p = 0.07 F(2,84) = 3.20, p = 0.05 | #1 (air v. ethanol) T2 p=0.026 #2 (air) T1,T2 p=0.012, T1,T3 p=0.000, T2,T3 p=0.036, T4,T6 p=0.000, T5,T6 p=0.009 #3 (ethanol) T1,T2 p=0.000, T1,T3 p=0.000 |
| 75 | Fig.S2Bv. Fig.S2G (latency) | | day*trials*group: χ2(2) = 28.95, p = 0.00, ε=0.66 F(1.33,55.76) = 3.91, p = 0.04 | #1 (air v. ethanol) day 1 p=0.010, T2 p=0.006, T3 p=0.053, T5 p=0.024 #2 (air) T1,T3 p=0.031, T2,T3 p=0.040, T4,T6 p=0.004, T5,T6 p=0.016 #3 (ethanol) T1,T2 p=0.000, T1,T3 p=0.000, T2,T3 p=0.016, T3,T6 p=0.005, T4,T5 p=0.002 |
| 76 | Fig.S2J. (distance) | | day*trials*group: χ2(2) = 6.09, p = 0.05 F(2,84) = 5.54, p = 0.01 | day 1 v day 2 p=0.008 #1 (air v. ethanol) T3 p=0.027 #2 (air) T1,T2 p=0.025 #3 (ethanol) T1,T2 p=0.014, T1,T3 p=0.000, T1,T4 p=0.000, T2,T5 p=0.034, T3,T6 p=0.008, T5,T6 p=0.05 |
| 77 | Fig.S2Cv. Fig.S2K. (time) | | trials: χ2(2) = 8.74, p = 0.01, ε=0.84 F(1.68,70.47) = 4.54, p = 0.02 | day 1 v. day 2 p=0.008, T1,T2 p=0.027, T1,T3 p=0.015 |
| 78 | Fig.S3Ei. (EtOH Absorption, full assay) | **Administration** (air n=9, 50% ethanol n=9) Absorption and metabolism measured 0 and 50min post-exposure respectively | F(1.05,16.78) = 17.067, p=0.001 | (air v. ethanol) 5min p = 0.01, 10min p = 0.005, 15 & 20min p = 0.002, 25 & 30min p = 0.001 |
| 79 | Fig.S3Fi (EtOH Metabolism, full assay) | | F(1.21,9.31) = 7.21, p=0.011 | (air v. ethanol) 20min p = 0.042, 25min p = 0.015, 30min p = 0.005. |
| | | | a.) Levene's Test for equality of variances b.) Independent Group t-test | NA |
| 80 | Fig.S3Eii. mM EtOH conc., (mM/Fly) | **Administration** (air n=9, 50% ethanol n=9) Absorption and metabolism measured 0 and 50min post-exposure respectively | F(16) = 0.763, p=0.395 t(16) = -3.603, p = 0.002 | NA |
| 81 | Fig.S3Fii. mM EtOH conc., (mM/Fly) | | F(16) = 0.147, p=0.706 t(16) = -1.214, p = 0.242 | |
| 82 | Fig.S3G. 50% ethanol absorption and metabolism | | F(16) = 5.215, p=0.036 t(16) = 3.976, p = 0.002 | |

Table 2f. Statistics Table

| | Figure | Data Structure, n-value | Statistical Test<br>1.) Mauchly's Test of Sphericity. If sphericity is violated (sig. < 0.05) then Greenhouse-Geisser correction (ε) applied to 2.) Repeated Measures Anova | Post Hoc<br>Bonferroni correction w/ pairwise comparisons |
|---|---|---|---|---|
| 83 | Fig S5A.<br>yaw | **All Data - Runway perframe features**<br>(air n=12, 50% ethanol n=12) z-scores for each behavior plotted in a heatmap | trials: χ2(2) = 0.22, p = 0.90<br>F(2,44) = 5.15, p = 0.01 | NA |
| 84 | angle2walll | | day*trials*group: χ2(2) = 4.66, p = 0.10<br>F(2,44) = 3.60, p = 0.04 | #1 (air v. ethanol) day 2 p=0.014, T1 p=0.015, T6 p=0.007 #2 (air) T1,T2 p=0.046, T1,T4 p=0.012, T2,T3 p=0.008, T3,T6 p=0.009 |
| 85 | phisideways | | day*trials*group: χ2(2) = 3.40, p = 0.18<br>F(2,44) = 1.70, p = 0.19 | NA |
| 86 | absdtheta | | trials: χ2(2) = 4.47, p = 0.11<br>F(2,44) = 68.43, p = 0.00 | T1,T2 p=0.000, T1,T3 p=0.000, T4,T5 p=0.000, T4,T6 p=0.000 |
| 87 | absdv_cor | | trials: χ2(2) = 3.95, p = 0.14<br>F(2,44) = 41.35, p = 0.00 | T1,T2 p=0.000, T1,T3 p=0.000, T4,T5 p=0.000, T4,T6 p=0.000 |
| 88 | signdtheta | | day*group: χ2(2) = 2.21, p = 0.33<br>F(1,22) = 6.96, p = 0.02 | #1 (air v. ethanol) day 1 p=0.03, T1 p=0.023 |
| 89 | ddist2wall | | trials*group: χ2(2) = 2.99, p = 0.22<br>F(2,44) = 4.02, p = 0.03 | #1 (air v. ethanol) day 2 p=0.036, T1 p=0.05 |
| 90 | absyaw | | trials: χ2(2) = 1.35, p = 0.51<br>F(2,44) = 8.67, p = 0.00 | T1,T3 p=0.003, T4,T5 p=0.022, T4,T6 p=0.008 |
| 91 | velmag_ctr | | trials: χ2(2) = 2.26, p = 0.32<br>F(2,44) = 37.01, p = 0.00 | T1,T2 p=0.000, T1,T3 p=0.000, T4,T5 p=0.000, T4,T6 p=0.000 |
| 92 | phi | | day*trials*group: χ2(2) = 3.50, p = 0.17<br>F(2,44) = 0.34, p = 0.71 | NA |
| 93 | flipdv_cor | | trials: χ2(2) = 0.02, p = 0.99<br>F(2,44) = 18.36, p = 0.00 | T1,T2 p=0.002, T1,T3 p=0.004, T4,T5 p=0.003, T4,T6 p=0.003 |
| 94 | dphi | | day*trials*group: χ2(2) = 1.76, p = 0.42<br>F(2,44) = 5.78, p = 0.01 | #1 (air v. ethanol) T3 p=0.001, T4 p=0.055 #2 (air) T2,T3 p=0.012 #3 (ethanol) T1,T3 p=0.05, T3,T6 p=0.004 |
| 95 | theta | | trials*group: χ2(2) = 8.67, p = 0.01, ε=0.75<br>F(1.50,32.88) = 5.10, p = 0.02 | #1 (air v. ethanol) T2 p=0.007 #2 (air) T1,T2 p=0.007, T2,T3 p=0.015 |
| 96 | dist2wall | | day*trials*group: χ2(2) = 5.24, p = 0.07<br>F(2,44) = 1.05, p = 0.36 | NA |
| 97 | angle2corner | | day*trials*group: χ2(2) = 8.65, p = 0.01, ε=0.75<br>F(1.50,32.90) = 2.46, p = 0.11 | NA |
| 98 | Fig S5A.<br>thrashing | **All Data - Runway classifier**<br>(air n=12, 50% ethanol n=12) z-scores for each behavior plotted in a heatmap | day*trials*group: χ2(2) = 16.02, p = 0.00, ε=0.65<br>F(1.30,28.69) = 3.01, p = 0.08 | NA |
| 99 | pacing | | day*trials*group: χ2(2) = 1.81, p = 0.41<br>F(2,44) = 0.06, p = 0.94 | NA |
| 100 | retreating | | day*trials*group: χ2(2) = 2.39, p = 0.30<br>F(2,44) = 0.50, p = 0.61 | NA |
| 101 | advancing | | day*trials*group: χ2(2) = 0.62, p = 0.73<br>F(2,44) = 0.42, p = 0.66 | NA |
| 102 | pausing | | days*trials: χ2(2) = 4.11, p = 0.13<br>F(2,44) = 3.20, p = 0.05 | T1,T2 p=0.011, T1,T3 p=0.027, T4,T5 p=0.003, T4,T6 p=0.030 |

**Additional Supplementary Item 3. Perframe feature descriptions.**

| Perframe Feature | | Description |
|---|---|---|
| absdv_cor | 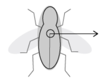 | sideways speed of the fly's center of rotation (\|dv_cor\|) (mm/s) [1 x (nframes -1)] |
| absdtheta | 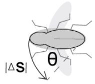  | angular speed (rad/s) (\|theta\|) [1 x (nframes -1)] |
| angle2corner_rect | 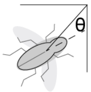  | angle to closest arena corner, relative to the fly's orientation (rad) [1 x nframes] |
| angle2wall_rect | 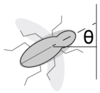  | angle to closest point on the arena wall, relative to the fly's orientation (rad) [1 x nframes] |
| absyaw | 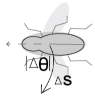  | absolute difference between velocity direction and orientation (rad) [1 x nframes] |
| ddist2wall_rect | 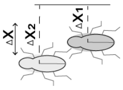  | change is distance to arena wall (mm/s) [1 x (nframes -1)] |
| dist2wall_rect | 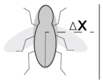  | distance to arena wall (mm) [1 x nframes] |
| dphi | 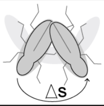  | change in velocity direction (rad/s) [1 x (nframes -1)] |
| flipdv_cor | 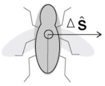  | sideways velocity of the fly's center of rotation, sign-normalized so that if the fly's orientation is turning right, then flipdv_cor is positive if the fly's center of rotation is also translating to the right (dv_cor * signdtheta) (mm/s) [1 x (nframes -1)] |
| phi | 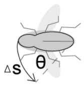  | velocity direction (rad) [1 x nframes] |
| phisideways | 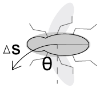  | difference between velocity direction and direction orthogonal to fly's orientation (range is [-pi/2,pi/2]) (rad) [1 x nframes] |
| signdtheta | 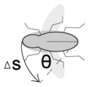  | boolean expressing whether change in orientation is positive or negative (no units) [1 x (nframes -1)] |
| theta | 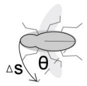  | angular velocity (rad/s) [1 x (nframes -1)] |
| velmag_ctr | 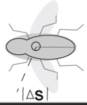  | speed (mm/s) [1 x (nframes -1)] |
| yaw | 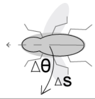  | difference between velocity direction and orientation (rad) [1 x nframes] |

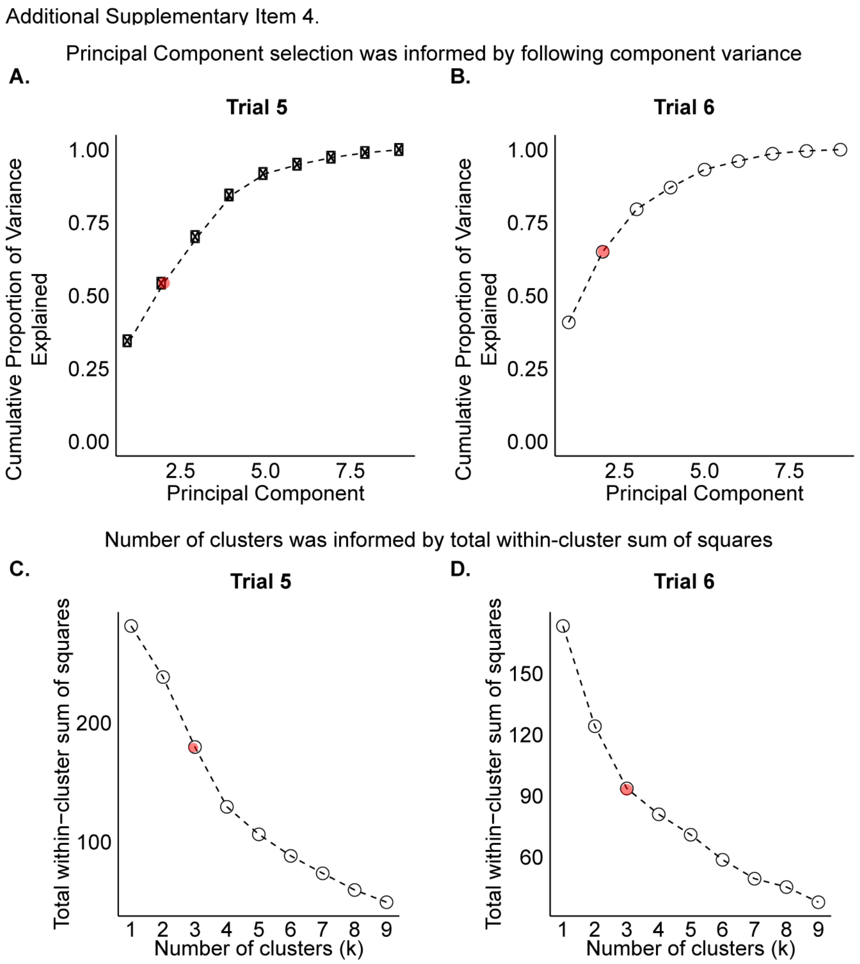


**Additional Supplementary Item 4. PCA component variance and total-within cluster sum of squares.** PCA plots were constructed for trial 5 and trial 6, where each fly was plotted as a function of 14 behaviors (time, latency, distance, angular velocity, advancing, pausing, pacing, thrashing, dist2wall, signdtheta, yaw, angle2corner, angle2wall, phisideways). Each trial's Euclidean distance matrix was projected onto its first 2 principal components. (**A-B**) Cumulative proportion of variance indicates that (A) 39% of trial 5 variance is represented by PC1 and PC2 [see ST4.] and (B) 44% of trial 6 variance is represented by PC1 and PC2. (**C-D**) Total within-cluster sum of squares indicates that number of clusters k=3 is sufficient to represent (**C**) trial 5 and (**D**) trial 6. (**E**) Principal Component Analysis component variance and variable loadings. PCA analysis was performed on 24 flies (air n = 12, 50% ethanol n = 12) for trial 5 and trial 6. Each trial's Euclidean distance matrix was projected onto its first 2 principal components. This table includes the standard deviation, proportion of variance, and cumulative proportion of variance of PC1-PC4 for trial 5 and trial 6, and the variable loadings of PC1 and PC2 for trial 5 and trial 6.

Additional Supplementary Item 4. (E) PCA component variance and variable loadings

**Component Variance**

| | Trial 5 | | | | Trial 6 | | | |
|---|---|---|---|---|---|---|---|---|
| | **PC1** | **PC2** | **PC3** | **PC4** | **PC1** | **PC2** | **PC3** | **PC4** |
| Standard deviation | 2.05 | 1.55 | 1.4 | 1.32 | 1.75 | 1.35 | 1.05 | 0.75 |
| Proportion of variance | 0.22 | 0.17 | 0.15 | 0.14 | 0.25 | 0.19 | 0.15 | 0.11 |
| Culumlative proportion | 0.22 | 0.39 | 0.54 | 0.68 | 0.25 | 0.44 | 0.59 | 0.7 |

**Variable loadings for PC1 and PC2**

| | Trial 5 | | Trial 6 | |
|---|---|---|---|---|
| | **PC1** | **PC2** | **PC1** | **PC2** |
| absyaw | 0.113 | -0.643 | 0.504 | -0.347 |
| advancing | 0.043 | 0.362 | -0.512 | 0.145 |
| angle2wall_rect | 0.118 | -0.466 | -0.226 | -0.523 |
| pausing | 0.215 | -0.195 | 0.384 | -0.328 |
| theta | -0.046 | -0.334 | 0.406 | 0.587 |
| velocity | -0.791 | -0.212 | 0.001 | 0.113 |
| absdtheta | -0.254 | 0.121 | -0.125 | 0.283 |
| latency | 0.397 | -0.094 | 0.310 | 0.192 |
| angular_velocity | 0.274 | 0.142 | 0.086 | -0.056 |
